# Alfalfa transcriptomic responses to the field pathobiome

**DOI:** 10.1101/2025.01.21.634162

**Authors:** Lev G. Nemchinov, Brian M. Irish, Sam Grinstead, Olga A. Postnikova

## Abstract

The pathobiome is a comprehensive biotic environment that includes a community of all disease-causing organisms within the plant, defining their mutual interactions and the resultant effect on plant health. We have previously reported on the diverse composition of the alfalfa pathobiome in the field production environment. We have also briefly addressed host reactions to a consortium of pathogenic microorganisms by profiling gene expression patterns shaped by the plant pathobiome in the field. In this study, we expand on the ‘field host genomics’ approach by surveying gene expression changes in visually healthy and diseased plants collected from different commercial alfalfa fields. In addition to offering insights into the genetic basis of host resistance to multi-pathogenic infections in natural ecosystems, this strategy can also help to identify plants with tolerant genotypes adapted to field pathobiome that could be used in breeding programs.

## Introduction

With the development of modern high-throughput sequencing approaches, the “one microbe - one disease” concept is being largely reevaluated and replaced with the principle of a “pathobiome”. A pathobiome is a comprehensive biotic environment that includes a diverse community of all disease-causing organisms within the plant (Vayssier-Taussat et al., 2014; Bass et al., 2019; Mannaa and Seo, 2021). While numerous individual pathogens and diseases they cause in alfalfa (*Medicago sativa* L.), the most extensively cultivated forage legume in the world, were described in detail, the concept and understanding of the crop’s pathobiome remain largely unexplored. We have recently demonstrated that alfalfa pathobiome represents a sophisticated habitat which encompasses at least 20-30 potential pathogens in individual plants (Nemchinov et al, 2023). In theory, such a complex multi-pathogenic environment may not only affect the behavior of all coinfecting organisms, their accumulation in the host, virulence, disease etiology and epidemiology, but also impact host fitness, prompting unique genetic responses to the collective pathobiome. These responses could be assessed for their roles in resistance to multiple biological stressors in the natural field environment rather than to a single pathogen under controlled experimental conditions, following the standard practice. To the best of our knowledge, this “field host genomics” approach is missing in the literature and has yet been undertaken in alfalfa or any other crop. Meanwhile, revealing genetic basis of host resistance to multi-pathogenic infections can improve our understanding of pathogenicity and further accelerate breeding programs.

In this work, we applied the proposed strategy to estimate host genetic responses to a collective pathobiome in commercial alfalfa fields. Since in the natural environment, all field plants are impacted by various organisms to some degree, it is not possible to obtain *de facto* uninfected control samples for transcriptomic analysis. To overcome this issue in experiments described herein, visually asymptomatic plants with healthy appearance were selected from the same fields where symptomatic/diseased plants were collected. While healthy-looking plants may contain various microbial communities of commensal, mutualistic or latent nature, organisms inhabiting them do not result in any symptoms and disease development with minimal to no damage to the host, if any (Sinclair, 1991). Importantly, these non-affected plants collected from the same fields as diseased samples, may potentially be considered tolerant or resistant to multi-pathogenic infections (Hull, 2014). Using modern transcriptomics tools combined with computational analyses, we assessed gene expression patterns in symptomless alfalfa plants and compared theses to samples displaying obvious symptomatology and disease damage. As a result of this work, genes and pathways involved in alfalfa responses to a diverse field pathobiome were identified and the genetic basis of resistance to multi-pathogenic infections was proposed.

### Materials and Methods Plant materials and sample collection

Samples were collected with owners’ permission from five commercial alfalfa fields in neighboring Benton and Yakima Counties, Washington State, USA during mid-July 2024 **(Table 1**). Five leaves, including all three leaflets and petiolules, were collected from each of five different asymptomatic and separately from five symptomatic plants of similar stage of development in five individual fields. The materials from five asymptomatic and separately from five symptomatic plants collected from each of the five fields were pooled for nucleic acid extraction to allow more efficient and comprehensive screening of a large population of plants (i.e. field). This resulted in two pooled samples from each field and 10 samples total that is, five biological replications for each sample type (asymptomatic and symptomatic). Asymptomatic samples were provisionally designated as “healthy” (H) and symptomatic “diseased” (D). Sampling was done in a zigzag or “W” pattern to include representative samples across the entire field. After collection, samples were immediately placed on dry ice and subsequently stored at −80C until extraction. Diseased plants displayed a variety of symptoms including yellowing, mosaic, mottling, stunting, chlorosis, local lesions, dwarfing, leaf spots, wilts, blights, anthracnose, crown and stem rot, decline, and stunting, while “healthy” plants did not exhibit any obvious symptoms (**Fig. 1**).

**Table 1.** Plant sample types and locations.

| Field № | Sample ID | Identifier | Area | County | Latitude/Longitude |
| --- | --- | --- | --- | --- | --- |
| 1 | H1 | Healthy | Prosser, WA | Benton | 46.253329, -119.730632 |
| 1 | D1 | Diseased | Prosser, WA | Benton | 46.253329, -119.730632 |
| 2 | H2 | Healthy | Prosser, WA | Benton | 46.243104, -119.758655 |
| 2 | D2 | Diseased | Prosser, WA | Benton | 46.243104, -119.758655 |
| 3 | H3 | Healthy | Prosser, WA | Benton | 46.227857, -119.791274 |
| 3 | D3 | Diseased | Prosser, WA | Benton | 46.227857, -119.791274 |
| 4 | H4 | Healthy | Grandview, WA | Yakima | 46.202759, -119.867621 |
| 4 | D4 | Diseased | Grandview, WA | Yakima | 46.202759, -119.867621 |
| 5 | H5 | Healthy | Grandview, WA | Benton | 46.221344, -119.863790 |
| 5 | D5 | Diseased | Grandview, WA | Benton | 46.221344, -119.863790 |

**Figure 1.**
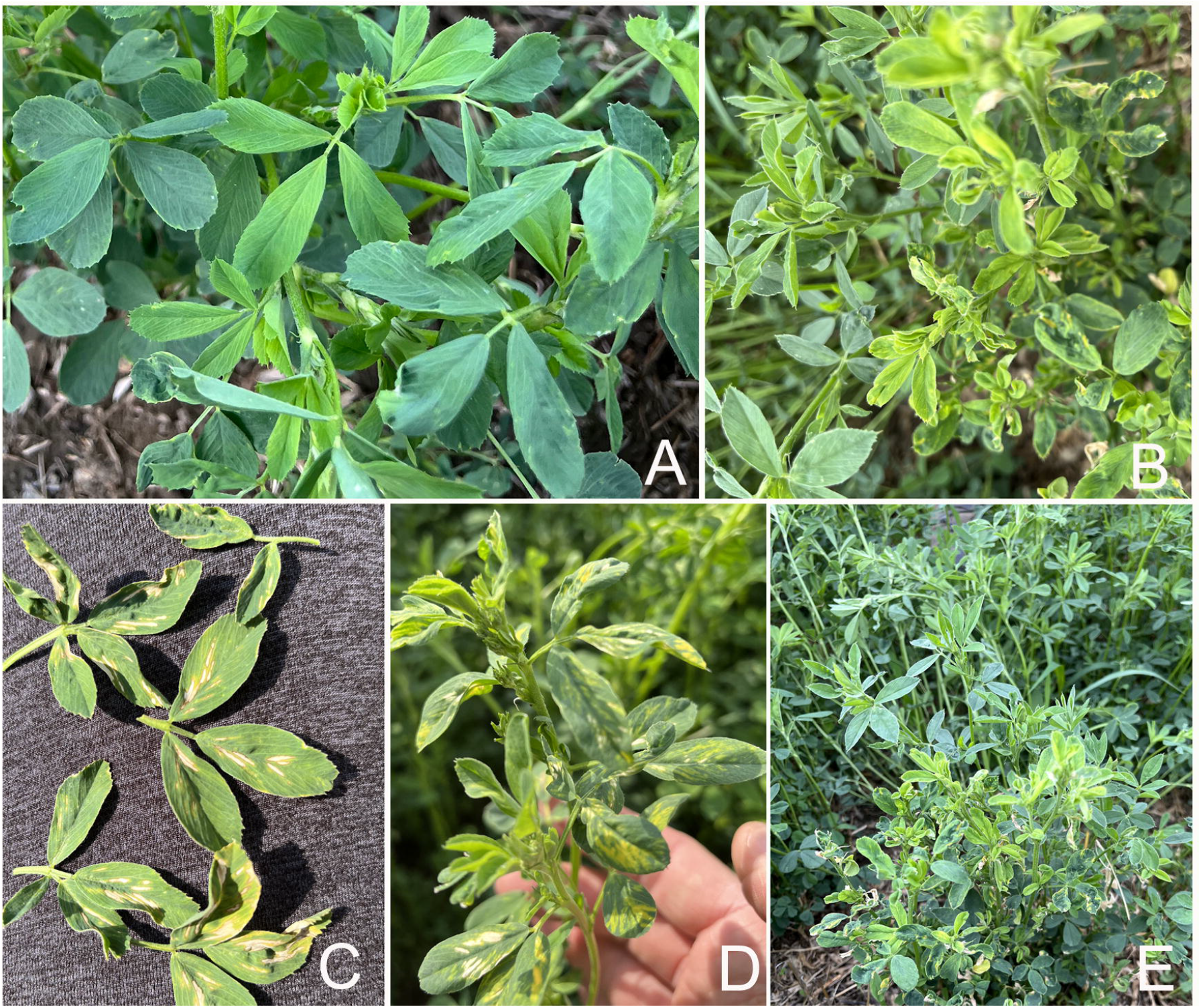
Phenotypes of asymptomatic (**A**) and symptomatic (**B, C, D, E**) alfalfa plants. **B**, leaf proliferation. **C**, leaf necrotic lesions. **D**, viral symptoms, likely AMV. **E**, shoot proliferation and stunting.

### Nucleic acid extraction

Pooled leaves from symptomatic and asymptomatic plants were pulverized in liquid nitrogen and used for RNA and DNA extractions. Total RNA extractions were performed with Qiagen’s RNeasy Mini Kit (50) according to manufacturer’s directions (Qiagen Inc., Germantown MD, USA), and genomic DNA extractions were done using Qiagen’s DNeasy PowerSoil Pro Kit (50) per the manufacturer’s instructions. In our hands, DNeasy PowerSoil Pro Kit worked better than any other kit for DNA extraction from plant tissues.

### RNA and Amplicon sequencing

Library preparation and Illumina RNA sequencing were performed by Novogene (Novogene Corp., Sacramento, CA USA). Messenger RNA was purified from total RNA using poly-T oligo-attached magnetic beads. The cDNA library was prepared using ABclonal Fast RNA-seq Lib Prep Kit V2 (ABclonal Technology, Wobum MA, USA). The library was checked with a Qubit and real-time PCR for quantification and a bioanalyzer for size distribution detection. Quantified libraries were sequenced on Illumina NovaSeq X Plus platform (PE150; 12 Gb or 40 million reads per sample).

The 16S and ITS (Internal Transcribed Spacer) library preparation for ribosomal RNA (rRNA) sequencing was carried out using an ABclonal Rapid Plus DNA Lib Prep Kit (Abclonal, Wobrun, MA USA). Libraries were sequenced on Illumina NovaSeq 6000 platform (PE250, 50K raw tags per sample). To discriminate from chloroplast DNA, primers targeted amplification of the DNA encoding V5-V7 region of the bacterial 16S rRNA gene: 799F, 5′ AACMGGATTAGATACCCKG 3′ and 1193R, 5′ ACGTCATCCCCACCTTCC 3′, with Illumina adapters. Primers used for amplification of the fungal internal transcribed spacer (ITS) region were ITS1F (F) 5′ CTTGGTCATTTAGAGGAAGTAA 3′ and ITS2R (R) 5′ GCTGCGTTCTTCATCGATGC 3′ (Nemchinov et al, 2023).

### Bioinformatics analysis

Bioinformatics analysis was performed both in-house and as a part of the RNA and Amplicon sequencing service provided by Novogene. Novogene’s workflow for bioinformatic analysis is shown in **Fig. S1**. Reads were assembled with StringTie (Pertea et al. 2015) and quantified with featureCounts (Liao et al. 2014). For differential analysis, DESeq2 and edgeR were used (Love et al. 2014; Robinson et at., 2010, respectively). Gene expression was estimated by the abundance of transcripts that mapped to the alfalfa reference genome (Chen et al., 2020) genome. FPKM values (Fragments Per Kilobase of transcript sequence per Millions base pairs sequenced) (Trapnell et al., 2010) were used for estimating gene expression levels, taking into consideration the effect of both sequencing depth and gene length on counting of fragments (Mortazavi et al., 2008) Alignment to the reference was performed with HISAT2 software (Mortazavi et al. 2008). For functional analyses, including GO (Gene Ontology) Enrichment, DO (Disease Ontology) Enrichment, and KEGG (Kyoto Encyclopedia of Genes and Genomes), clusterProfiler was used (Yu et al. 2012). Gene Set Enrichment Analysis (GSEA) was performed using GSEA software (https://www.gsea-msigdb.org/gsea/index.jsp). Principal component analysis (PCA) was used to evaluate intra-and -intergroup differences (Jolliffe, I.T. and Jorge Cadima, J. 2016). The protein-protein interaction network was constructed by searching protein interaction database STRING (https://string-db.org). SNP/InDel Analysis was performed by snpEff tool (Cingolani et al. 2012). QIIME analysis (Quantitative Insights Into Microbial Ecology), (Caporaso et al. 2010) was used to identify fungal and bacterial communities in 16S and ITS sequencing data. OTU (Operational Taxonomic Units) clustering was employed for taxonomic annotations. OTUs were analyzed for relative abundance, taxonomic abundance, common and unique groups (Venn diagram, and Flower diagrams), and phylogenetic relationships. Alpha and Beta Diversity analyses were performed to analyze composition of microbial communities in different samples.

Bioinformatics pipeline for virus identification was used essentially as described in Nemchinov et al. (2023).

## Results

### 1. Field pathobiome of symptomatic and asymptomatic alfalfa plants

The complexity of alfalfa’s field pathobiome was evaluated with an emphasis on the composition of viral, bacterial, and fungal communities.

#### 1.1 Viruses

Total number of plant viral reads for all diseased plants was 7,710,052 and for all healthy-appearing plants 19,407.753. These numbers were dominated by pea streak virus (PeSV), 7,333,345 and 19,028,984 reads, in asymptomatic and symptomatic plants, respectively. Although PeSV is usually symptomless in alfalfa, it is transmissible to susceptible legume hosts, like pea and lentil crops (Larsen, 2015). All other plant viruses excluding pea streak virus, had 376,707 and 378,769 reads, respectively. Therefore, both asymptomatic and symptomatic plants were infected to approximately similar extent with the same viruses (**Fig. 2**). The predominant species were PeSV, Medicago sativa alphapartitivirus 1, alfalfa mosaic virus (AMV), Snake River alfalfa virus (SRAV), alfalfa rhabdoviruses, alfalfa latent virus (ALV), and bean leafroll virus (BLRV) – all present in asymptomatic and symptomatic plants. As some of these viruses, especially AMV, may cause distinct symptomatology, one could suggest that healthy-appearing plants collected from the same field as diseased plants, exhibit tolerance to the infection possibly due to genetic heterogeneity, resistance to aphids or absence of seed transmission.

**Figure 2.**
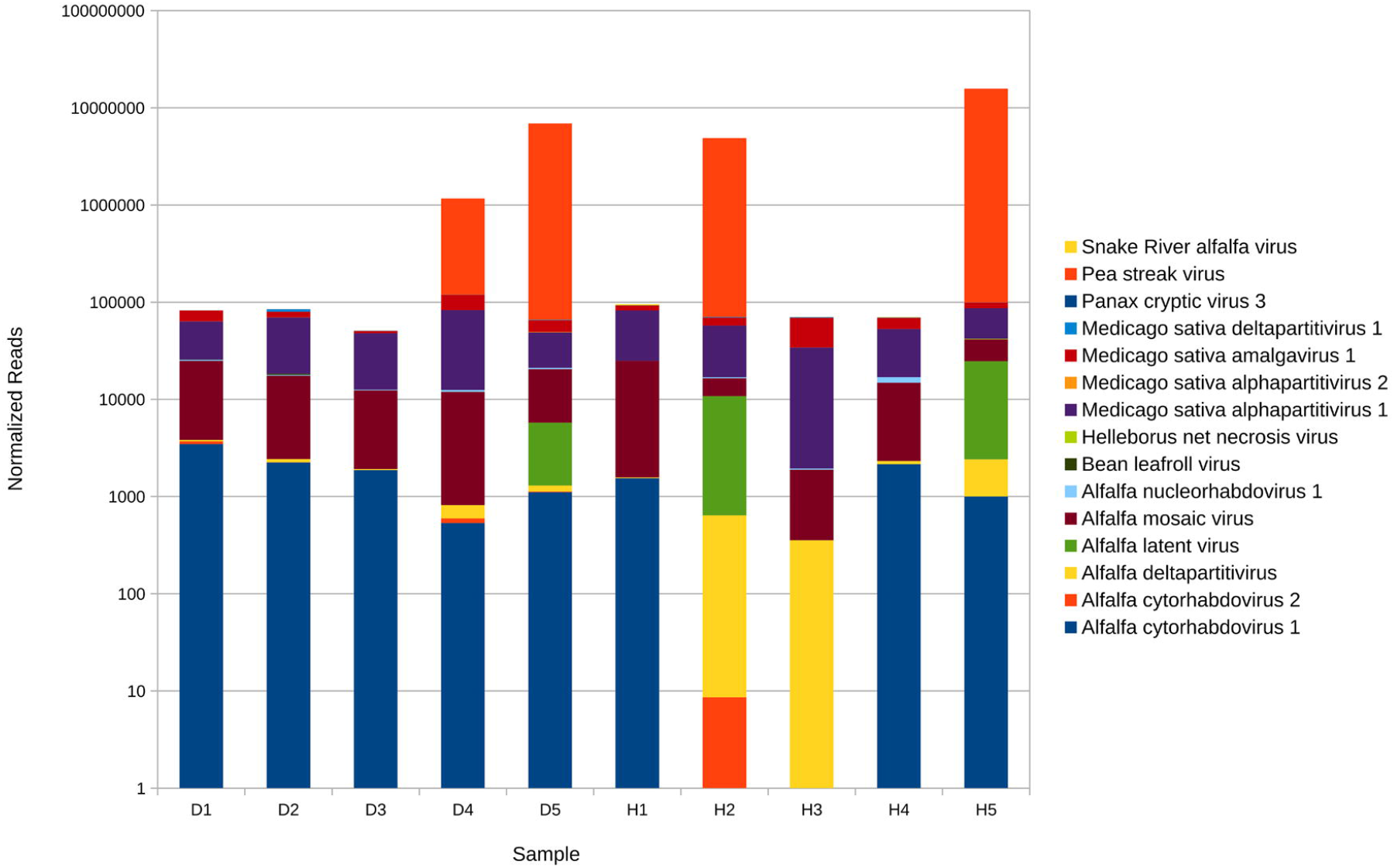
Composition and relative abundance of the alfalfa (*Medicago sativa* L.) virome in 50 symptomatic (D) and asymptomatic (H) plant samples based on the results of RNA-sequencing.

#### 2.2 Bacteria

As a result of the 16S rRNA sequencing, 1,222,165 clean bacterial reads were obtained that were annotated to OTUs followed by their classification at the level of kingdom, phyla, class, order, family, genus and species (**Table 2**). Bacterial species identified across the two groups are shown in **Table S1**. Composition and diversity of the identified OTUs were assessed by relative and taxonomic abundance analyses, and by alpha and beta diversity index values of amplicon sequence data.

**Table 2.** Clean bacterial reads classified into different taxonomic ranks.

| **Sample** | **Kingdom** | **Phylum** | **Class** | **Order** | **Family** | **Genus** | **Species** |
| --- | --- | --- | --- | --- | --- | --- | --- |
| H1.2 | 37187 | 37152 | 37136 | 36700 | 35953 | 32436 | 9240 |
| D1.2 | 49026 | 48999 | 48991 | 48904 | 48107 | 45284 | 18075 |
| H2.2 | 47419 | 47376 | 47364 | 47048 | 39847 | 36892 | 14421 |
| D2.2 | 69503 | 69496 | 69496 | 69485 | 66863 | 66218 | 38713 |
| H3.2 | 50920 | 50735 | 50735 | 50438 | 47667 | 42306 | 18142 |
| D3.2 | 46404 | 46380 | 46378 | 46241 | 42700 | 38274 | 15377 |
| H4.2 | 69213 | 69198 | 69198 | 69171 | 69090 | 68530 | 65188 |
| D4.2 | 83197 | 83184 | 83182 | 83174 | 72883 | 71774 | 40054 |
| H5.2 | 60917 | 60868 | 60840 | 60805 | 55641 | 52425 | 19903 |
| D5.2 | 47000 | 46986 | 46983 | 46953 | 43978 | 37559 | 15632 |

Predominant relative abundance was registered for the populations of bacteria belonging to phyla *Actinobacteriota, Proteobacteria*, and *Bacillota* (*Firmicutes*) (**Fig. 3**). While most actinobacteria and some firmicutes are beneficial to plants, many species in the phylum *Proteobacteria* are pathogenic. On average, distribution of relative abundance was comparable between symptomatic and asymptomatic plants. The taxonomic abundance cluster heatmap showed that diseased samples had a noticeable presence of several pathogenic species (*Pseudomonas viridiflava, Agrobacterium rubi, Acidovorax* sp.) (**Fig. S2**) absent in asymptomatic plants, which were often infected with known beneficial bacteria. Nevertheless, similar to our previous study on the alfalfa pathobiome (Nemchinov et al. 2023), the occurrence of pathogenic bacteria in symptomatic alfalfa field samples was not extensive.

**Figure 3.**
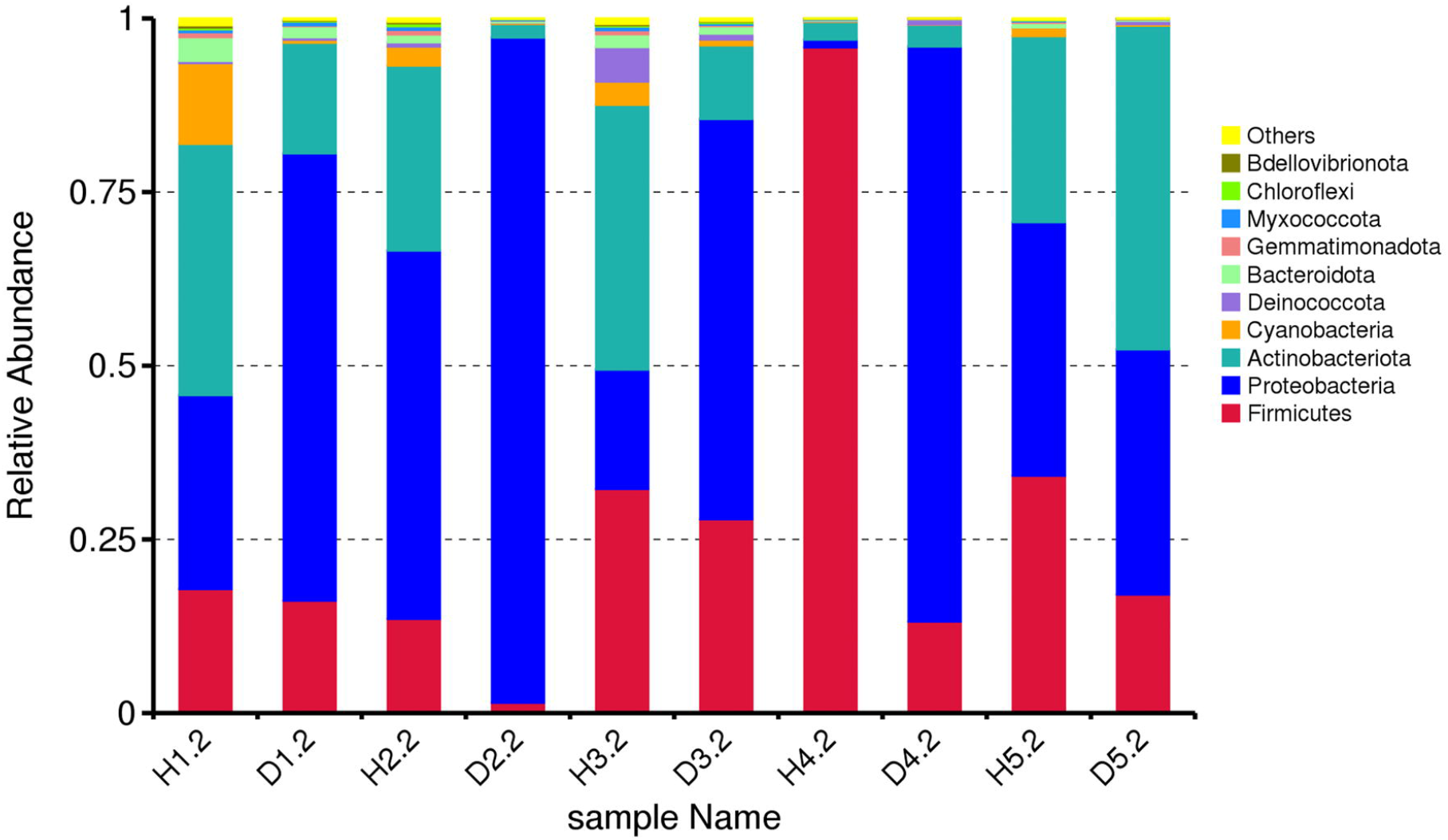
Composition and relative abundance of the alfalfa (*Medicago sativa* L.) bacteriome at the phyla level based on the results of 16S V5-V7 amplicon sequencing.

Alpha diversity analysis performed to assess richness of microbial communities in each group, showed greater diversity of bacterial species in asymptomatic plants (**Fig. S3**). Meanwhile, the beta-diversity heatmap indicated that dissimilarity coefficients between bacterial communities of symptomatic and asymptomatic plants originated from the same fields were generally low (**Fig. S4**), thus indicating an overall low species diversity.

Therefore, analogously to the virome, 16S rRNA sequencing demonstrated that bacteriome of asymptomatic and symptomatic alfalfa plants in the field was comparable.

#### 1.2. Fungi

In total, 972,315 clean combined reads were obtained by the ITS sequencing. Sequences were annotated to the level of kingdom, phyla, class, order, family, genus and species (**Table 3**). Species identified in all samples are shown in **Supplementary Table 1**. Relative abundance registered at the species level showed the presence of pathogenic *Stemphylium sp*., *Ascochyta medicaginicola, Alternaria* sp., *Cladosporium herbarum, Leptosphaerulina* sp., *Fusarium* sp., and others in symptomatic and asymptomatic plants (**Fig. 4**). All samples contained reads of *Amphinema* sp., corticioid fungi in the family *Atheliaceae*, known as wood-rotting species. The taxonomic cluster heatmap showed the abundance of the species in each sample, confirming the output of relative abundance distribution (**Fig. S5**). Alpha diversity analysis indicated a greater diversity of observed species in asymptomatic samples (**Fig. S6**), while beta diversity analysis suggested, with few exceptions, an overall low level of dissimilarities between samples originating from the same fields (**Fig. S7**). Once again, although diseased plants were infected with several known fungal pathogens, plants displaying no visual symptoms were not pathogen-free.

**Table 3.**
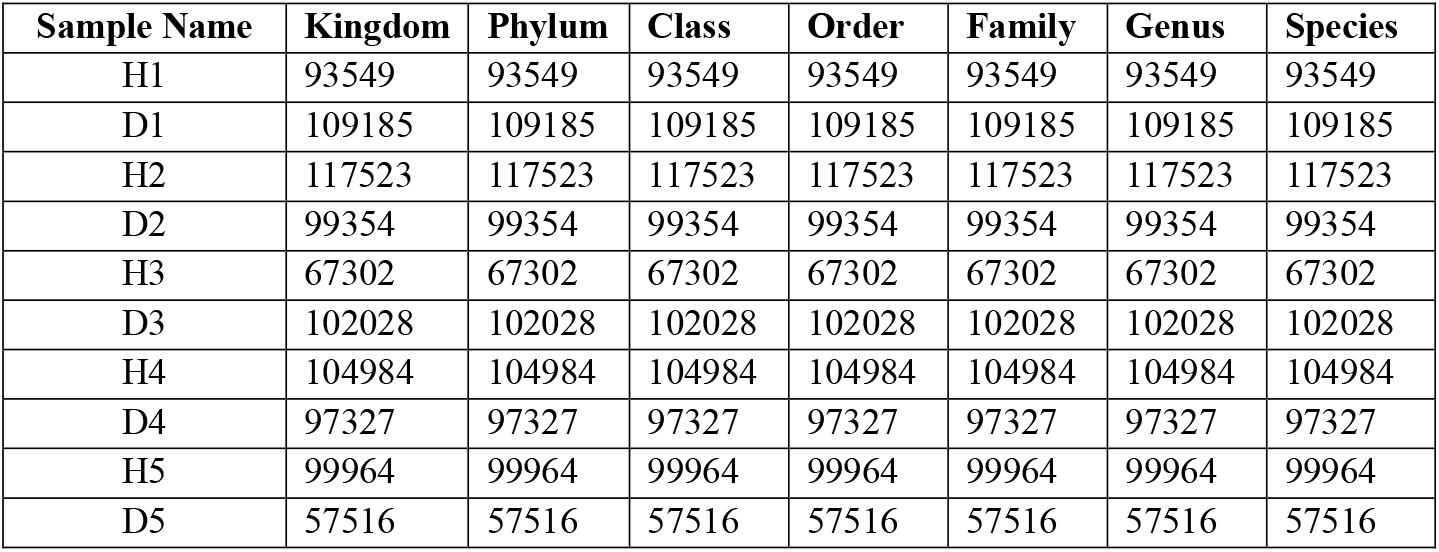
Clean fungal reads classified into different taxonomic ranks.

**Figure 4.**
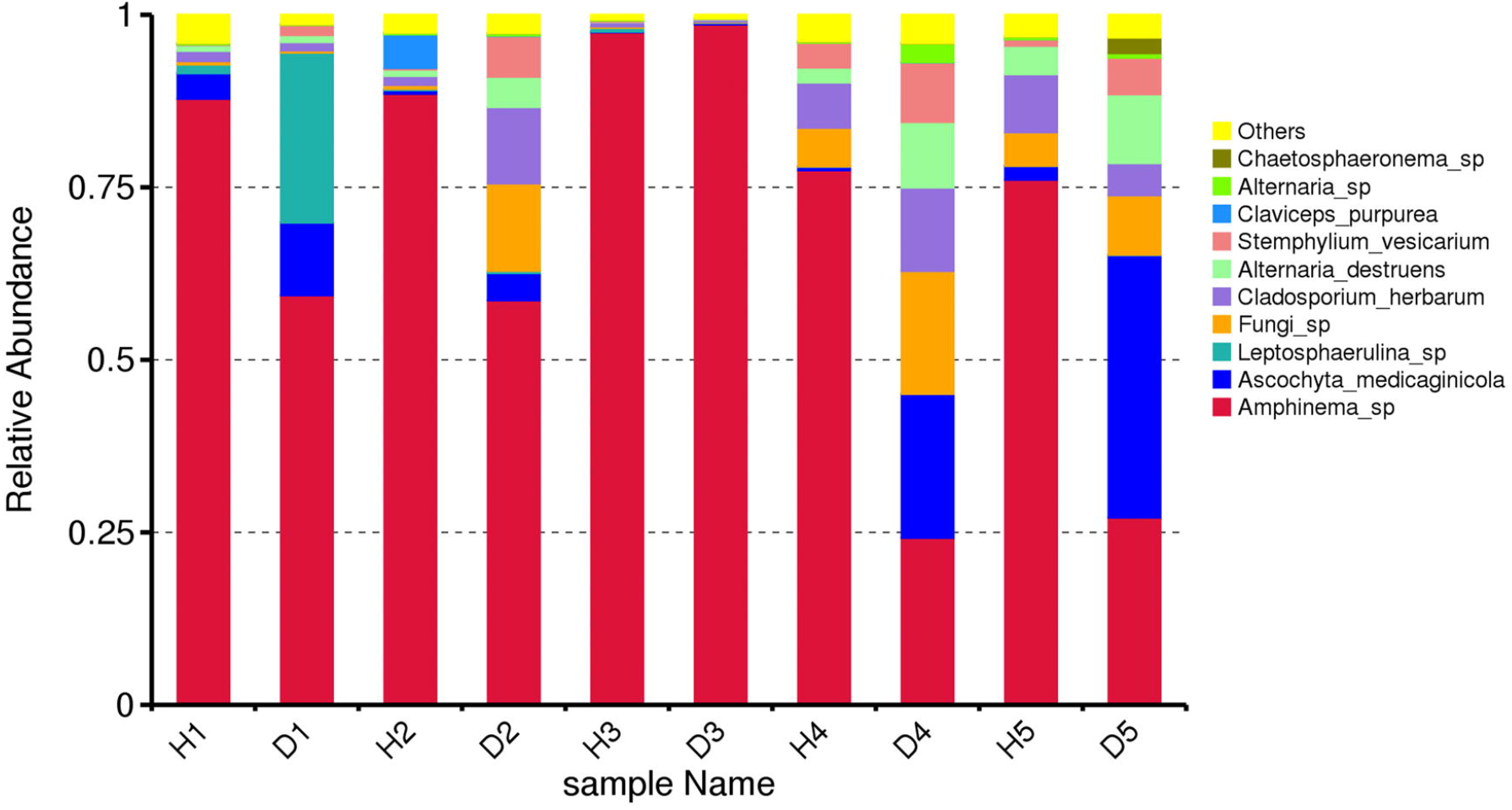
Composition and relative abundance of fungal species in alfalfa (*Medicago sativa* L.). field samples based on the results of ITS1-2 amplicon sequencing.

### 2. Gene expression in symptomatic vs asymptomatic alfalfa plants

#### 2.1 Metrics of RNA-Seq Data

Metrics of RNA-seq data are shown in **Table 4**. A total of 837,870,186 clean paired-end reads were generated from 10 cDNA libraries, thus averaging 83,787,019 paired-end or 41,893,509.3 single-reads per library. The alignment rate for each library ranged between 69% and ~ 87% mapped to the reference alfalfa genome (Chen et al. 2020). The RNA sequencing error rate was low 0.01 (1 in 10,000 bases), indicating high-quality sequencing acceptable for the experiment. Overall, the data obtained were sufficient for gene expression profiling.

**Table 4.** Metrics of RNA-seq data.

| Sample | Raw reads | Clean reads | Error rate | Total map to genome |
| --- | --- | --- | --- | --- |
| H1 | 110398188 | 104593218 | 0.01 | 88307654(84.43%) |
| D1 | 84350094 | 77925090 | 0.01 | 65403007(83.93%) |
| H2 | 95274306 | 88743738 | 0.01 | 70321916(79.24%) |
| D2 | 85298434 | 79717748 | 0.01 | 65901676(82.67%) |
| H3 | 85367348 | 79914866 | 0.01 | 67642954(84.64%) |
| D3 | 80221118 | 75485700 | 0.01 | 63847887(84.58%) |
| H4 | 93298768 | 89186996 | 0.01 | 77547028(86.95%) |
| D4 | 94300706 | 88395878 | 0.01 | 73903545(83.61%) |
| H5 | 83551536 | 77985652 | 0.01 | 54318556(69.65%) |
| D5 | 80042856 | 75921300 | 0.01 | 57685071(75.98%) |
| **Total reads** | 892103354 | 837870186 |  |  |

#### 2.2 Differentially expressed genes (DEGs) in symptomatic vs asymptomatic alfalfa plants

Two groups were assembled to analyze differential expression in symptomatic vs asymptomatic alfalfa phenotypes: one included all “healthy” samples collected from five different fields (H1-H5), and the other – all diseased samples (D1-D5). The first group was designated ‘A’ for asymptomatic and the second group ‘S’, for symptomatic phenotypes. A summary of DEGs stats between the S and A groups is shown in **Table 5** and their overall distribution, statistical significance and fold changes is presented in **Fig. 5**.

**Table 5.**
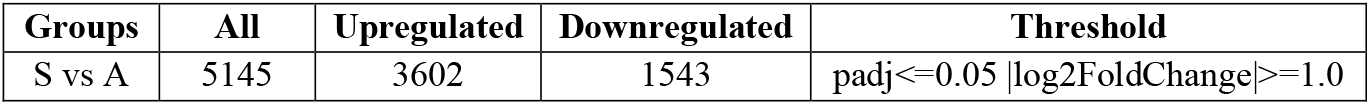
DEG summary stats between ‘S’ and ‘A’ phenotypes.

**Figure 5.**
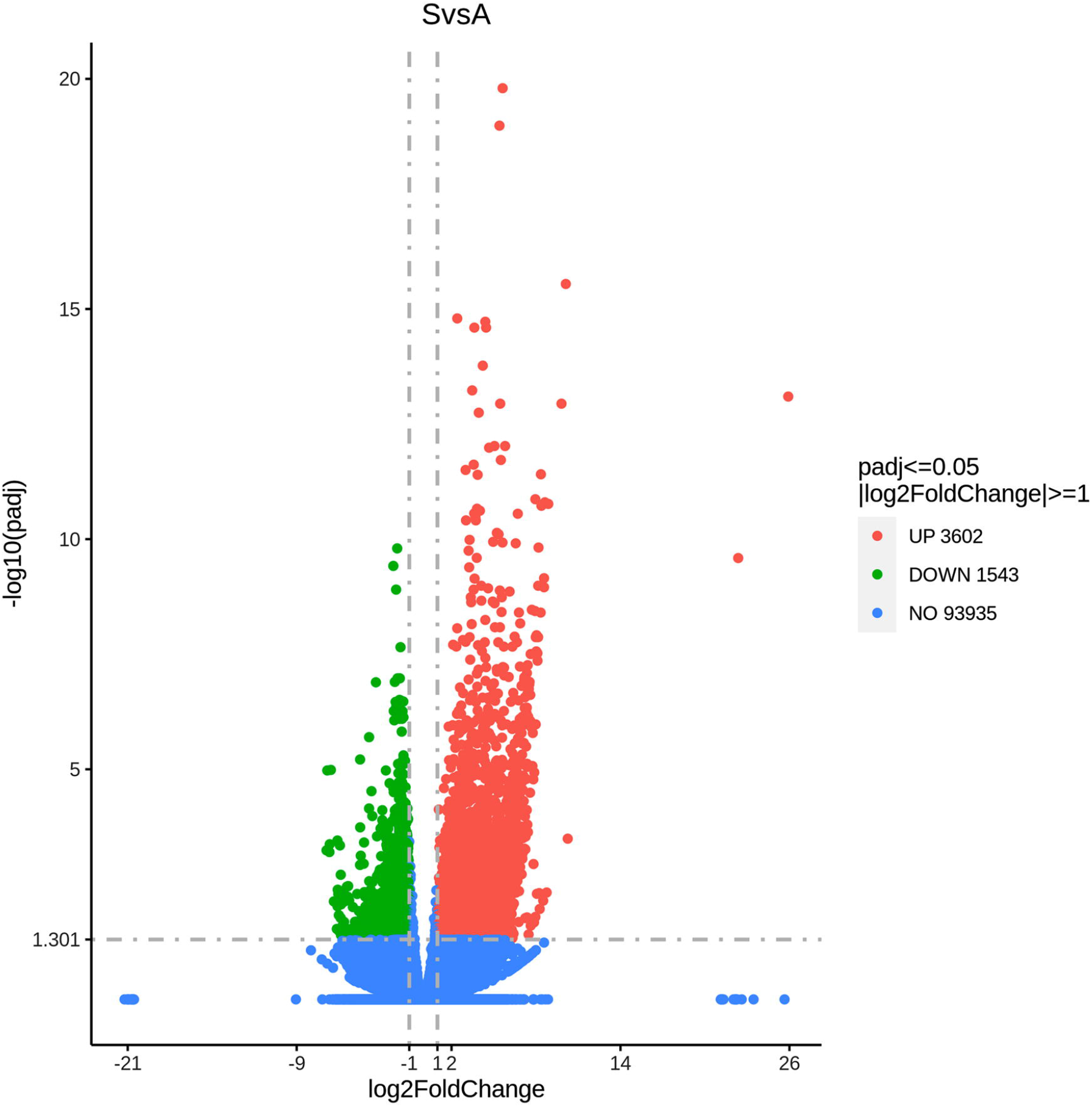
Volcano plot showing the overall distribution of differentially expressed genes. The x-axis shows the fold change in gene expression between different samples, and the y-axis shows the statistical significance of the differences. Red dots represent up-regulation genes and green dots represent down-regulation genes. The blue dashed line indicates the threshold line for differential gene screening criteria.

The total number of upregulated DEGs in symptomatic plants was significantly higher, indicating overall higher gene activity and mRNA production levels in diseased plants. Genes involved in RNA degradation (e.g. MS_gene91185, exosome complex component RRP43); proteosome components (e.g. MS_gene058875, 26S proteasome non-ATPase regulatory subunit 2); protein export (MS_gene0874; ATPase GET3B); disease resistance (e.g. MS_gene44944, MS_gene87339, etc.) and many other processes were upregulated in the diseased phenotypes suggesting vigorous anti-pathogenic responses are taking place. All individual DEGs, along with their expression values, gene IDs, and annotations are shown in **Tables S2, S4 and S4**. Cluster analysis was caried out to learn if genes with similar expression patterns can be grouped. Three groups of genes depicted expression changes between diseased and “healthy” plants: genes involved in photosynthetic processes, which were normally downregulated in diseased samples (**Fig. S8**); genes encoding heat shock proteins upregulated in diseased plants; and defense- and pathogenesis-related genes that had elevated expression in the symptomatic plants.

The most highly expressed genes in both phenotypes, according to the normalized read counts, were those involved in photosynthetic reactions (**Tables S2, S3 and S4**). Genes encoding ribulose-1,5-bisphosphate carboxylase/oxygenase (rubisco), that catalyzes fixation of carbon from atmospheric CO2 (Bathellier et al. 2020) and participates in photorespiration, were expressed at higher levels in asymptomatic phenotypes. Via the photorespiratory pathway, rubisco can mitigate oxidative stress, down-regulating production of reactive oxygen species in the chloroplast during immune response and protect photosystems from photodamage (Jiang et al. 2023; Voss et al. 2013).

Oxygen-evolving enhancer proteins (PSBO), a critical part of photosynthetic complex important for thylakoid architecture and involved in host-pathogen interactions (Kong et al. 2014) and light-harvesting complex proteins, essential for plant photosynthetic machinery (Tanaka et al. 2010), were also highly expressed in asymptomatic phenotype vs. symptomatic phenotype.

Other known key regulators of plant immune response associated with chloroplasts, such as calcium-sensing receptor (CAS) mediating restriction of bacterial growth, gene encoding thylakoid formation protein (ThF1), mediating both PTI and ETI responses (Kachroo et al. 2021), and light-harvesting complex-like proteins, contributing to pathogen resistance and protection of chloroplasts from oxidative stress (Liu et al. 2019; Hey and Grimm, 2018), were expressed at higher levels in A-phenotype.

There were many silent or uniquely expressed genes (non-expressed or expressed at a very low level with near-zero read counts) in both phenotypes. When a gene is silent in A- and expressed in S-phenotype, this points to its activation due to the host-pathogen interactions, while when the opposite happens, it may indicate its role in mechanisms of resistance to field pathobiome.

The top 10 uniquely expressed genes upregulated in the ‘S’ and silent in the ‘A’ phenotype are shown in **Table 6**. Among them were those encoding Kunitz trypsin inhibitor involved in defense against pests and herbivores (Bonturi et a. 2022); heat shock proteins, including HSP90, mediating stress signal transduction; laccase (Lac 7) that plays important roles in plant defenses (Yu et al. 2021); VITVI Agamous-like MADS-box protein, a transcriptional regulator involved in stress responses (Castelán-Muñoz et al. 2019), and others. These genes were also on the top of the list of highly upregulated genes, thus emphasizing their vital roles in alfalfa interactions with the field pathobiome (Xu et al. 2012).

**Table 6.** Top ten uniquely expressed upregulated genes in ‘S’ phenotype.

| Gene ID | log2FoldChange | pvalue | padj | gene chr | Gene Description |
| --- | --- | --- | --- | --- | --- |
| MS_gene50908 | 25.91497945 | 1.10E-17 | 7.98E-14 | chr3.3 | [Q41015]PIP21 PEA Kunitz-type trypsin inhibitor-like 1 protein OS=Pisum sativum |
| MS_gene39508 | 22.36125335 | 1.52E-13 | 2.58E-10 | chr4.2 | [P27880]HSP12 MEDSA 18.2 kDa class I heat shock protein OS=Medicago sativa |
| MS_gene04350 | 10.25462041 | 3.47E-06 | 0.0003224 | chr3.4 | [P55737]HS902 ARATH Heat shock protein 90-2 OS=Arabidopsis thaliana |
| MS_gene048082 | 10.1199187 | 1.18E-20 | 2.86E-16 | chr4.4 | [Q9SR40]LAC7 ARATH Laccase-7 OS=Arabidopsis thaliana |
| MS_gene38363 | 9.810193692 | 1.88E-17 | 1.14E-13 | chr3.2 | [Q9SND9]Y3028 ARATH Uncharacterized acetyltransferase OS=Arabidopsis thaliana |
| MS_gene47682 | 8.872085208 | 5.67E-15 | 1.72E-11 | chr5.3 | [P24076]BGIA MOMCH Glu S.griseus protease inhibitor OS=Momordica charantia |
| MS_gene49154 | 8.758466235 | 0.0001264 | 0.0047591 | chr1.3 | [P51819]HSP83 IPONI Heat shock protein 83 OS=Ipomoea nil |
| MS_gene013756 | 8.632169259 | 5.01E-15 | 1.58E-11 | chr3.3 | [Q0HA25]MADS9 VITVI Agamous-like MADS-box protein MADS9 OS=Vitis vinifera |
| MS_gene04024 | 8.572115155 | 4.47E-13 | 7.06E-10 | chr3.3 | [Q01807]LEC2 MEDTR Truncated lectin 2 OS=Medicago truncatula |
| MS_gene011597 | 8.57085468 | 7.64E-13 | 1.11E-09 | chr8.1 | [Q9M7K4]POT5 ARATH Potassium transporter 5 OS=Arabidopsis thaliana |

Among the top 10 genes silent in the ‘S’ and expressed in the ‘A’ phenotype were haloacid dehalogenase-like hydrolase involved in stress responses (Zan et al. 2023); plasma membrane ATPase, participating in plant-microbe interactions (Elmore and Coaker, 2011); 40S ribosomal protein S3a, multifunctional protein regulating DNA repair, apoptosis, and plant innate immune response to bacterial infection (Gao and Hartwidge, 2011); DNA-binding domain of WRKY transcription factor, known for their role in plant immunity (**Table 7**), (Pandey and, Somssich, 2009), and others.

**Table 7.** Top ten uniquely expressed genes upregulated in ‘A’ phenotype.

| Gene_ID | log2FoldChange | pvalue | padj | gene_chr | Gene_Description |
| --- | --- | --- | --- | --- | --- |
| MS_gene97260 | -6.8757669 | 7.62E-06 | 0.0005771 | chr6.4 | Medicago truncatula uncharacterized LOC25495897 |
| MS_gene20930 | -6.8240864 | 4.38E-08 | 1.06E-05 | chr1.3 | \|PF13419: Haloacid dehalogenase-like hydrolase |
| MS_gene068502 | -6.6725796 | 8.60E-06 | 0.0006287 | chr4.2 | \|Q7XPY2\|PMA1_ORYSJ Plasma membrane ATPase OS=Oryza sativa subsp. japonica |
| MS_gene067994 | -6.6491031 | 5.01E-06 | 0.0004256 | chr8.3 | Hypothetical protein MtrunA17_Chr7g0215391 [Medicago truncatula] |
| MS_gene21834 | -6.569530738 | 4.22E-08 | 1.03E-05 | chr5.2 | \|Q8GTE3\|RS3A_CICAR 40S ribosomal protein S3a OS=Cicer arietinum |
| MS_gene87925 | -6.1429706 | 0.0016532 | 0.0294615 | chr6.1 | \|B8BJ39\|KPYC1_ORYSI Pyruvate kinase 1, cytosolic OS=Oryza sativa subsp. indica |
| MS_gene72202 | -6.0943342 | 0.0003328 | 0.0094798 | chr5.1 | \|PF03106: WRKY DNA -binding domain |
| MS_gene99515 | -6.0812176 | 0.0001038 | 0.0041409 | chr1.3 | \|Q6E593\|BEBT1_PETHY Benzyl alcohol O-benzoyltransferase OS=Petunia hybrida |
| MS_gene016372 | -6.0392219 | 0.0001403 | 0.0051221 | chr8.2 | \|PF00400: WD domain, G-beta repeat |
| MS_gene045088 | -6.0070489 | 0.0018772 | 0.0321011 | chr3.4 | \|Q02920\|NO70_SOYBN Early nodulin-70 OS=Glycine max |

Application of GO analysis to identify differentially expressed genes belonging to biological processes (BP), molecular functions (MF), and cellular components (CC) aspects, resulted in three clearly over-represented terms: photosynthesis (BP), thylakoid (CC), and protein heterodimerization activity (MF) (**Fig. 6)**. While the first two confirm essential vulnerability of photosynthetic reactions and structural elements to biotic stressors, the third term shows the importance of protein complexes formation in central parts of plant pathogenesis, including regulation of defense responses (Zhang et al. 2012).

**Figure 6.**
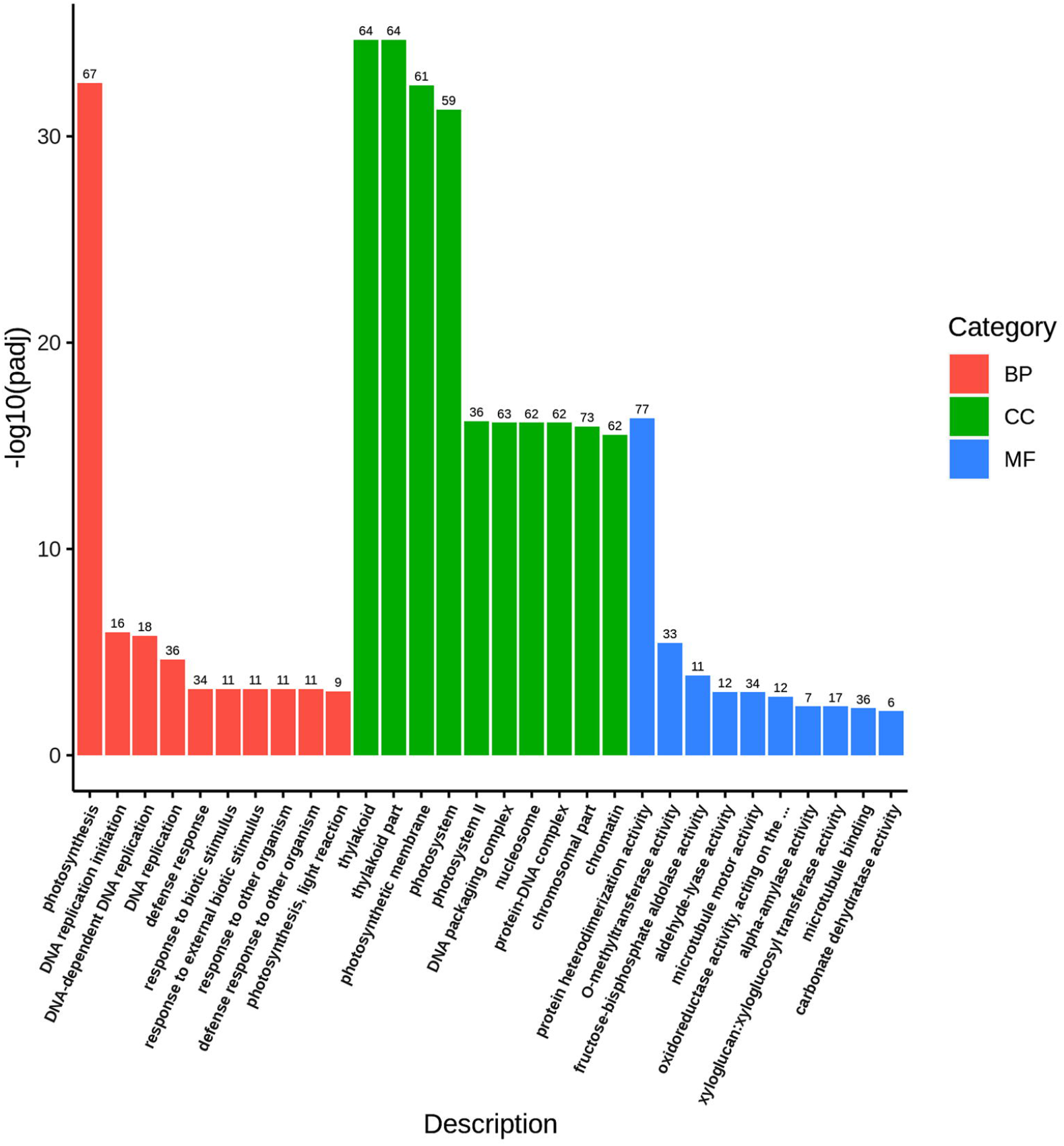
GO enrichment analysis histogram showing overrepresented terms. The abscissa represents a GO term, and the ordinate is GO term’s significance of enrichment expressed as - log10(padj). Different colors represent different functional categories.

GO enrichment analysis to identify overrepresented categories among upregulated genes showed that those mainly belonged to DNA replication processes and responses to stress, while enrichment of downregulated genes in the ‘S’ phenotype identified over-representation of photosynthesis-related genes and genes involved in energy metabolism (**Tables S5 and S6**). KEGG analysis, in general, confirmed these observations: key genes involved in photosynthesis were downregulated in the ‘S’ phenotype as compared to the ‘A’ phenotype (**Fig S9)**.

To further interpret gene expression data by estimating correlation between gene sets and the defined biological states/phenotypes (‘A’ and ‘S’), GSEA test was performed. GSEA showed 89 upregulated gene sets in the ‘S’ phenotype, of which 49 sets were significantly enriched at FDR < 25% and 23 sets were significantly enriched at nominal <1% (**Table S7**). In the ‘A’ phenotype, GSEA revealed 49 sets, of which 37 were significantly enriched at FDR < 25% and 20 gene sets were significantly enriched at nominal <1% (**Table S8**).

When using KEGG analysis, the top 10 enriched sets in phenotype ‘S’ were related to DNA replication, transporter proteins, chromatin remodeling, ribosome biogenesis, and recombination, while the top 10 enriched sets in phenotype ‘A’ were mostly associated with photosynthesis (**Table S7 and S8**). A heat map of the top 50 DEGs in each phenotype and ranked gene list correlation profile are shown in **Fig. S10**.

PCA analysis of the gene expression value (FPKM) was performed to evaluate intergroup and intragroup differences. PCA showed that samples of different phenotypes were clearly dispersed, while samples within each phenotype were generally clustered, indicating diverse expression patterns between symptomatic and asymptomatic plants and an analogous pattern within each group (**Fig. 7)**.

**Figure 7.**
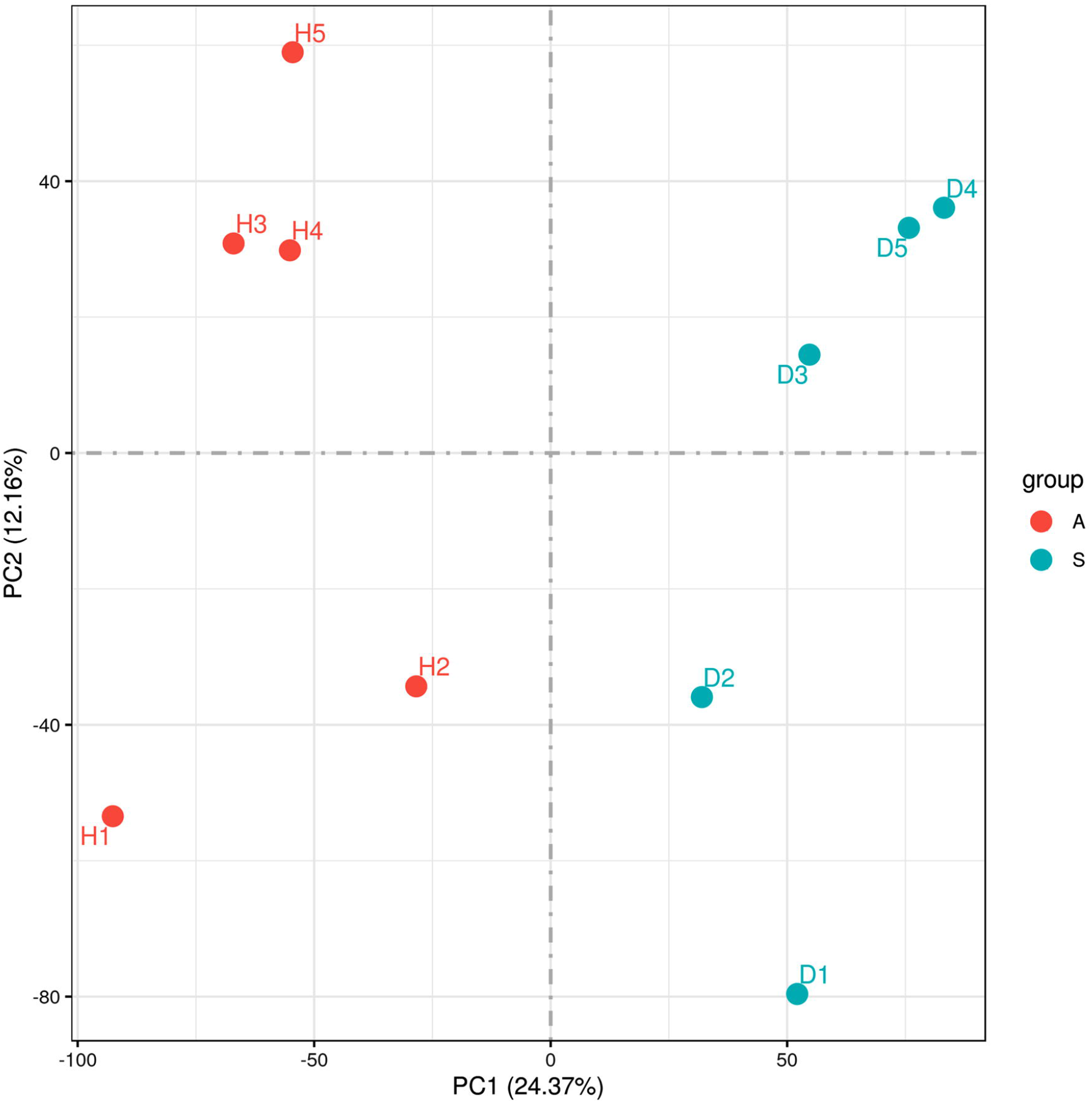
Principal component analysis (PCA) showing intergroup and intragroup differences. The PCA was performed based on the gene expression value (FPKM) of all samples.

Based on the analyses described above, two short list of genes were compiled including 1) genes upregulated in the symptomatic phenotype and likely involved in broad alfalfa responses to the collective pathobiome and 2) genes highly expressed in asymptomatic phenotype and potentially responsible for alfalfa tolerance to the collective pathobiome (**Table 8**).

**Table 8.**
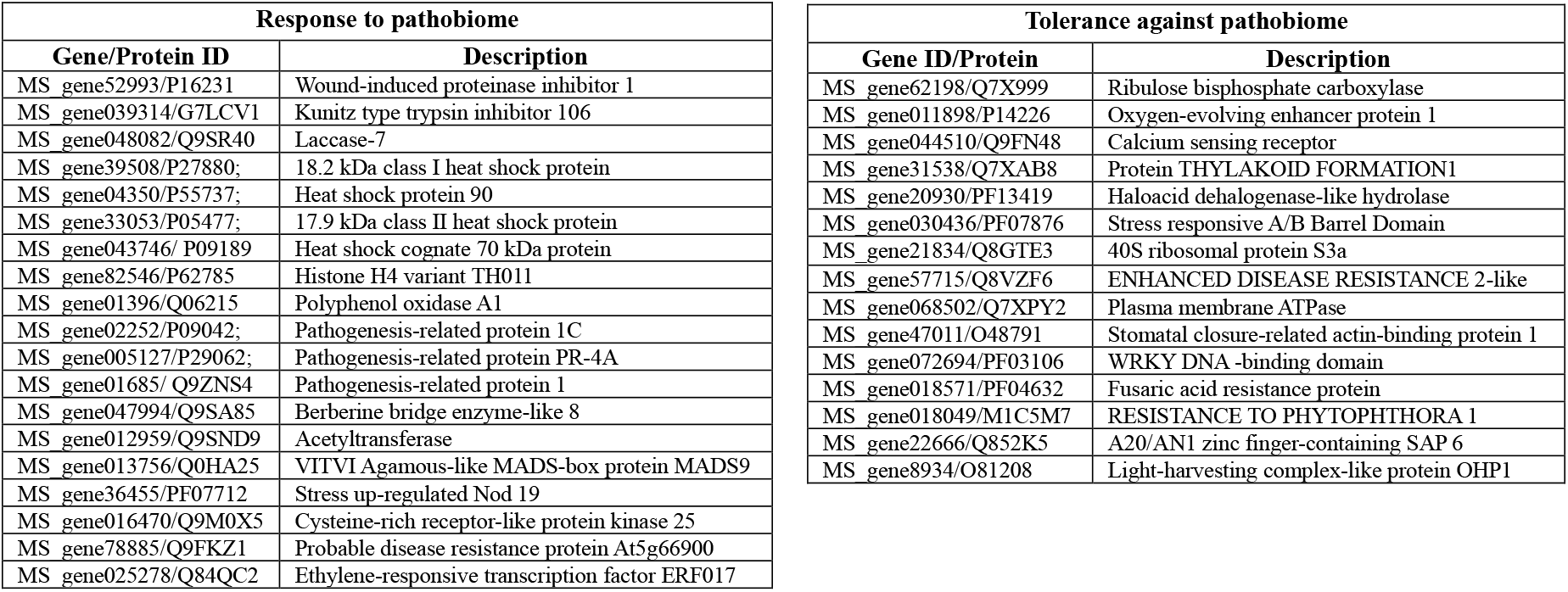
Short list of genes likely involved in alfalfa responses to the collective pathobiome.

#### 2.3 Interactions between proteins encoded by differentially expressed genes

To further understand reasons behind tolerance to multi-pathogenic infections, we analyzed protein-protein interactions (PPI) in asymptomatic plants. The physical interactions between proteins, although transient and dynamic, can affect many biological processes including plant defense and stress responses (Zhang et al. 2010; Struck et al. 2018).

PPI analysis showed that many proteins encoded by differentially expressed genes upregulated in symptomless plants (i.e. downregulated in diseased plants) and clustered together in the network, belong to the GO terms associated with photosynthesis (**Fig. S11)**. This implies that the proposed physical contacts between proteins are specific and have a particular biological meaning, which in this case would presumably be involvement in host-pathogen interactions and immune response (**Fig. S11)**; **Table S9**). Among the largest group of ~25 closely interacting proteins were light-harvesting chlorophyll a-b binding proteins, involved in in abscisic acid signaling (Liu et al. 2013); Photosystem II reaction center W protein, one of the central members of photosynthetic machinery (Rhee et al. 1998); Photosystem I reaction center subunit psaK, regulating light harvesting under stress (Amunts et al. 2009); Photosystem I reaction center subunit IV, essential for photosynthesis, and others (**Fig. S11**).

## Discussion

In this work, we applied a novel “field host genomics” approach toward the study of alfalfa transcriptomic responses to a collective field pathobiome represented by viral, bacterial, and fungal organisms. The approach aimed at the evaluation of genetic responses to a collective pathobiome in the natural field environment, rather than to a single pathogen under controlled experimental conditions. Allegedly, this strategy can offer insights into the genetic basis of host resistance to multi-pathogenic infections in natural ecosystems and help to identify plants with tolerant genotypes adapted to field pathobiome that could be used in breeding programs.

Identified pathogenic species previously reported as pathogenic on alfalfa included AMV, PeSV, BLRV, *Pseudomonas viridiflavia, Agrobacterium rubi, Acidovorax* sp., *Alternaria* sp., *Stemphylium* sp., *Ascochyta medicaginicola, Cladosporium herbarum, Claviceps purpurea* and other microorganisms. Plants were also infected with cryptic and not yet fully characterized viruses, beneficial and non-pathogenic bacterial and fungal species.

Notably, the microbiome of asymptomatic and symptomatic alfalfa field plants was found to be comparable, rather than distinct, which was hypothesized based on the presence or absence of the disease symptoms. This fact suggests that healthy-appearing plants may exhibit tolerance to the multi-pathogenic infections and are able to reduce the effect of the pathobiome on their overall fitness (Pagan and Garcia-Arenal, 2018). However, judging from the sequencing data, pathogens’ multiplication process was not limited in “healthy” plants, implying that resistance mechanisms (as opposed to tolerance) are less likely to play a pivotal role in lack of symptom expression.

Various reasons can potentially contribute to alfalfa tolerance to the pathobiome, considering that asymptomatic and symptomatic plants were collected from the same fields seeded with the same varieties, and growing under the same environment:

1. Tolerance observed in asymptomatic plants appears to be under genetic control, as they had uniquely expressed genes silent in symptomatic plants, and the number of DEGs and their expression level differed from the diseased phenotype.
2. Tolerant plants maintained at the upregulated level specific cellular processes, classified into enriched GO categories *photosynthesis, thylakoid, photosynthetic membrane, and oxidoreductase activity*. In other words, robust response of chloroplasts to multi-pathogenic infection may be critical for the establishment of tolerance as defense against microbial attack required energy provided by photosynthesis (Serrano et al. 2016). Chloroplasts play essential roles in plant defense reactions (Kachroo et al. 2021), and the majority of the disease symptoms observed in the field directly affected chloroplasts. Elevated expression of the genes encoding chloroplastic proteins in the ‘A’ phenotype can be associated not only with the absence of the disease symptomatology (i.e. undamaged chloroplasts), but also with effective immune response to pathogens attack that is, increased tolerance to the field pathobiome. This conclusion is supported by analysis of PPI in the asymptomatic plants.

Increased photosynthetic activity in tolerant plants resulted mainly from overexpression of genes encoding proteins catalyzing carbon fixation (e.g. MS_gene62198), structural components of chloroplasts (e.g. MS_gene31538) and proteins of Photosystems I and II (e.g. MS_gene03163 and MS_gene75549).

In addition to genes involved in photosynthesis, genes encoding proteins localized in chloroplasts and known to modulate defense responses were upregulated in tolerant plants, such as calcium-sensing receptor (MS_gene044510), thylakoid formation 1 protein (MS_gene31538), and light-harvesting complex-like protein OHP1 (MS_gene89341).

Genes not involved in photosynthesis also participated in tolerant response. Among those were HAD, regulating phosphate homeostasis during phosphorus (Pi) deficiency stress (Du et al. 2021); 40S ribosomal protein S3a with many roles including host-pathogen interactions (Gao and Hardwidge, 2011); disease resistance proteins; plasma membrane ATPase participating in plant immune responses (Elmore and Coaker, 2011), WRKY TFs, known for their role in plant immunity (Pandey and Somssich, 2009).

A simplified diagram, illustrating contribution of different pathways toward alfalfa tolerance to the field pathobiome is shown in **Fig. 8**. While combination of all relevant factors is important, chloroplasts appear to play a major role in promoting alfalfa’s ability to reduce the effect of multi-pathogenic infections on the plant fitness. Knowing genes controlling this important and likely polygenic trait can provide an alternative to host resistance-based management practices of plant diseases (Jeger, 2022).

**Figure 8.**
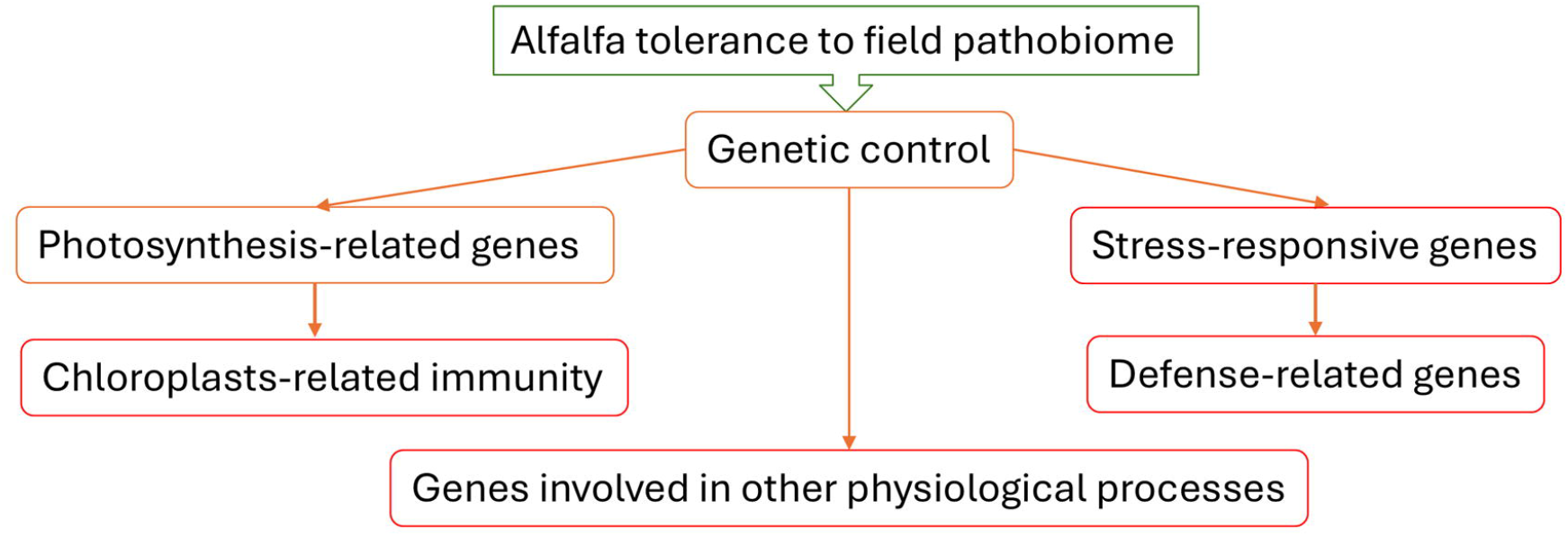
A simplified diagram, illustrating contribution of different pathways toward alfalfa tolerance to the field pathobiome.

## Acknowledgments

This study was supported by the United States Department of Agriculture, the Agricultural Research Service [CRIS numbers 8042-21000-300-000D (LN) and 2090-21000-026-000-D (BI)].

## Author contributions

LN: concept, data analysis, and first draft of the manuscript. BI: survey, sample collection, and evaluation. SG: bioinformatics and data analysis. OP: wet lab and data analysis. All authors contributed to the editing of the manuscript and approved it for publication.

## Data availability statement

The datasets have been submitted to the NCBI’s Sequence Read Archive (SRA) under the Submission ID: SUB14892281 and the BioProject accession ID: PRJNA1191001. They will be publicly available once the paper is confirmed for publication.

## Supplementary Tables

1. **Table S1**. Bacterial and fungal species identified in all samples.
2. **Table S2**. DEGs identified in ‘S’ and ‘A’ phenotypes.
3. **Table S3**. Upregulated DEGs identified in ‘S’ phenotype
4. **Table S4**. Downregulated DEGs identified in ‘S’ phenotype.
5. **Table S5**. Overrepresented GO categories among upregulated genes of phenotype ‘S’.
6. **Table S6**. Overrepresented GO categories among downregulated genes of phenotype ‘S’
7. **Table S7**. Gene sets enriched in phenotype ‘A’.
8. **Table S8**. Gene sets enriched in phenotype ‘S’
9. **Table S9**. Interactions between proteins encoded by genes downregulated in phenotype ‘S’ (upregulated in phenotype ‘A’).

## Supplementary Figures

**Figure S1.** Workflow of the bioinformatics analysis.

**Figure S2.** Taxonomic abundance cluster heatmap of bacterial species. The heatmap shows whether the samples with similar processing are clustered or not, while the similarity and differences between the samples can also be observed.

**Figure S3.** Alpha diversity of microbial communities in each group. The horizontal axis represents the groups, while the vertical axis represents the corresponding alpha diversity index value. Group G1.2, five asymptomatic plants; group G2.2, five symptomatic plants.

**Figure S4.** Beta diversity heatmap. The numbers in grids are the dissimilarity coefficient between samples. Two numbers in the same grid represent weighted and unweighted Unifrac distance, respectively.

**Figure S5.** Taxonomic cluster heatmap of fungal species.

**Figure S6.** Alpha diversity of the observed fungal species in each group. Group G1 composed of five asymptomatic plants; group G2– of five symptomatic plants.

**Figure S7.** Beta diversity heatmap showing diversity of the fungal communities between different sample. The numbers in grids are the dissimilarity coefficient between samples. Two numbers in the same grid represent weighted and unweighted Unifrac distance, respectively.

**Figure S8.** Heatmap depicting differentially expressed genes in both phenotypes.

**Figure S9.** KEGG pathway depicting downregulation of photosynthesis-related genes in symptomatic phenotype.

**Figure S10.** Heat map depicting top 50 DEGs in each phenotype.

**Figure S11.** Protein-protein interaction (PPI) network. The network was constructed by searching protein interaction database STRING (https://string-db.org) and visualized using Cytoscape software v.3.10.3. For GO enrichment, all connected nodes in the main network were selected and STRING functional enrichment option applied. **A**, Overrepresented GO terms are shown in green color. **B**, Non-redundant GO terms. **C**, Genes encoding interacting proteins (**Table S9**).

