## Supplementary figures and images for "Alfalfa transcriptomic responses to the field pathobiome"

### Fig.S1

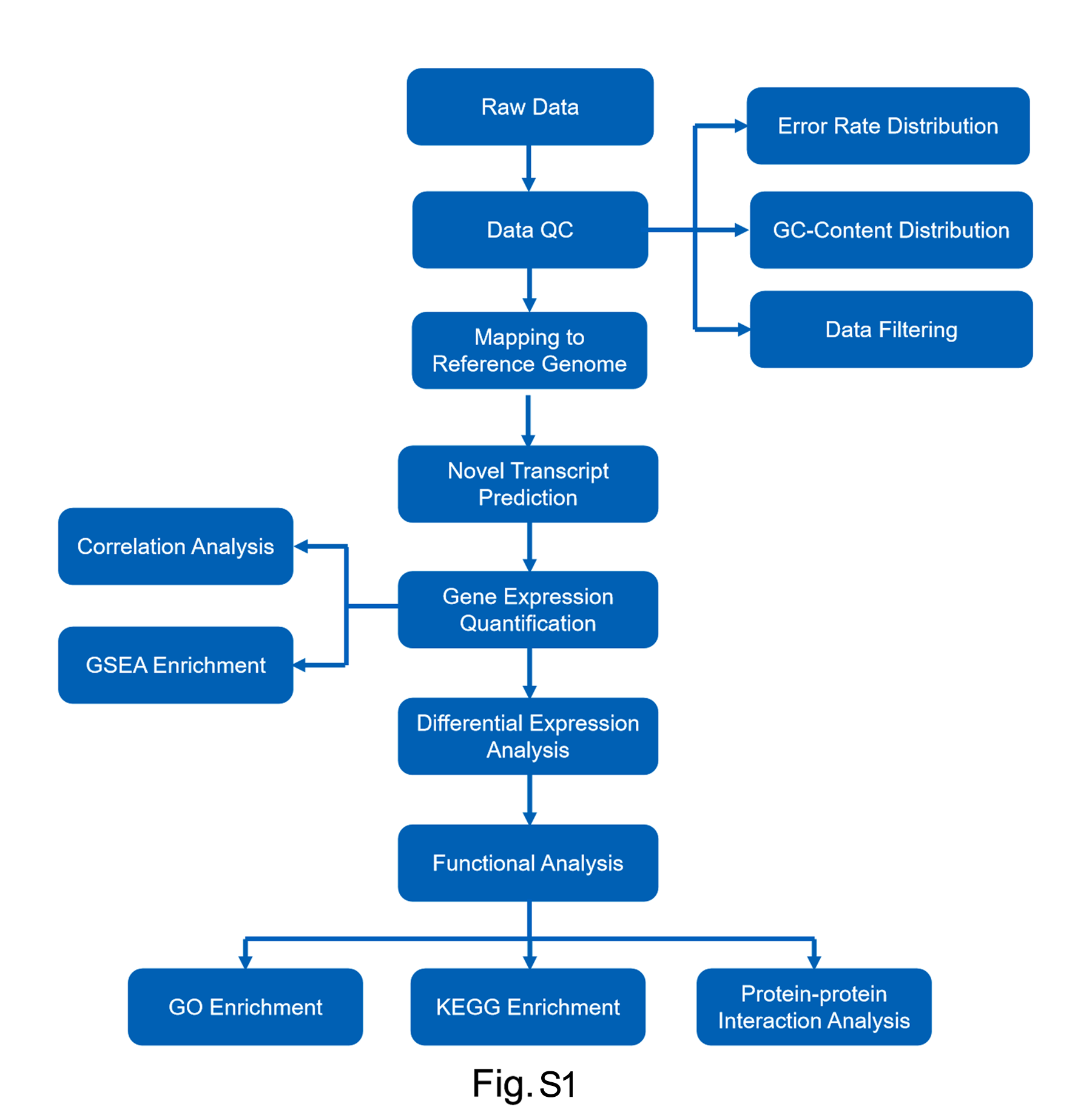

### Fig.S2

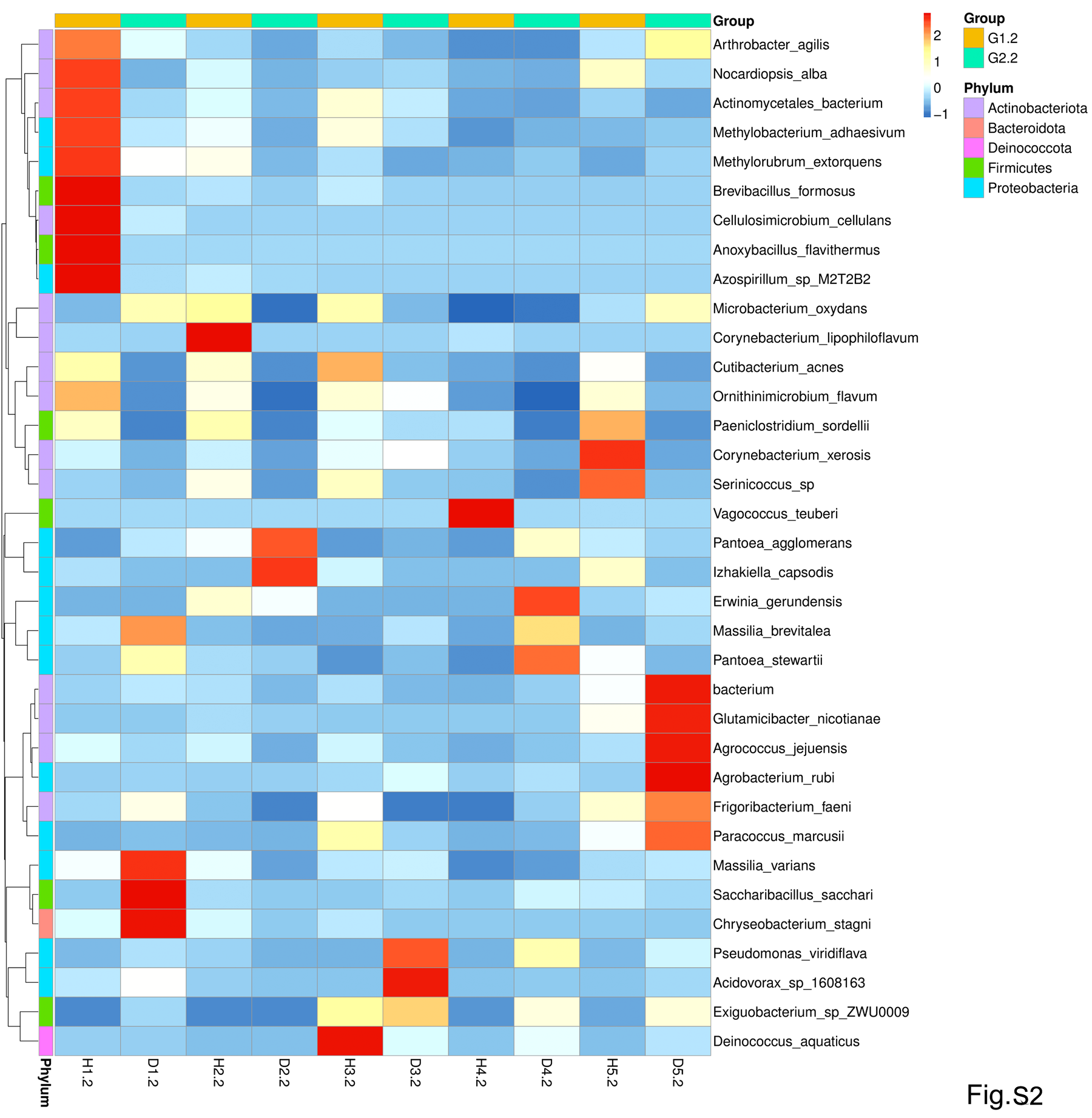

### Fig.S3

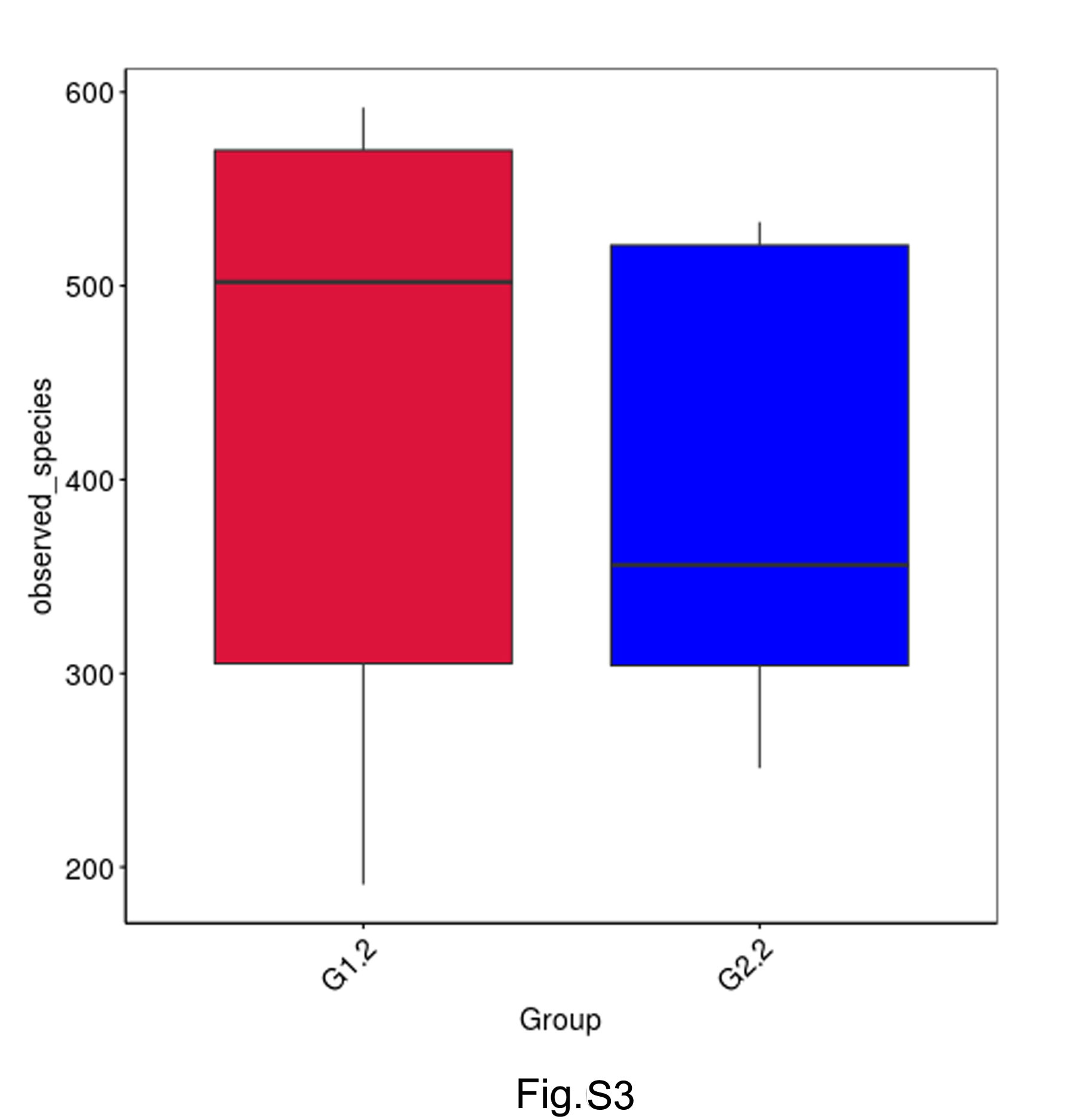

### Fig.S4

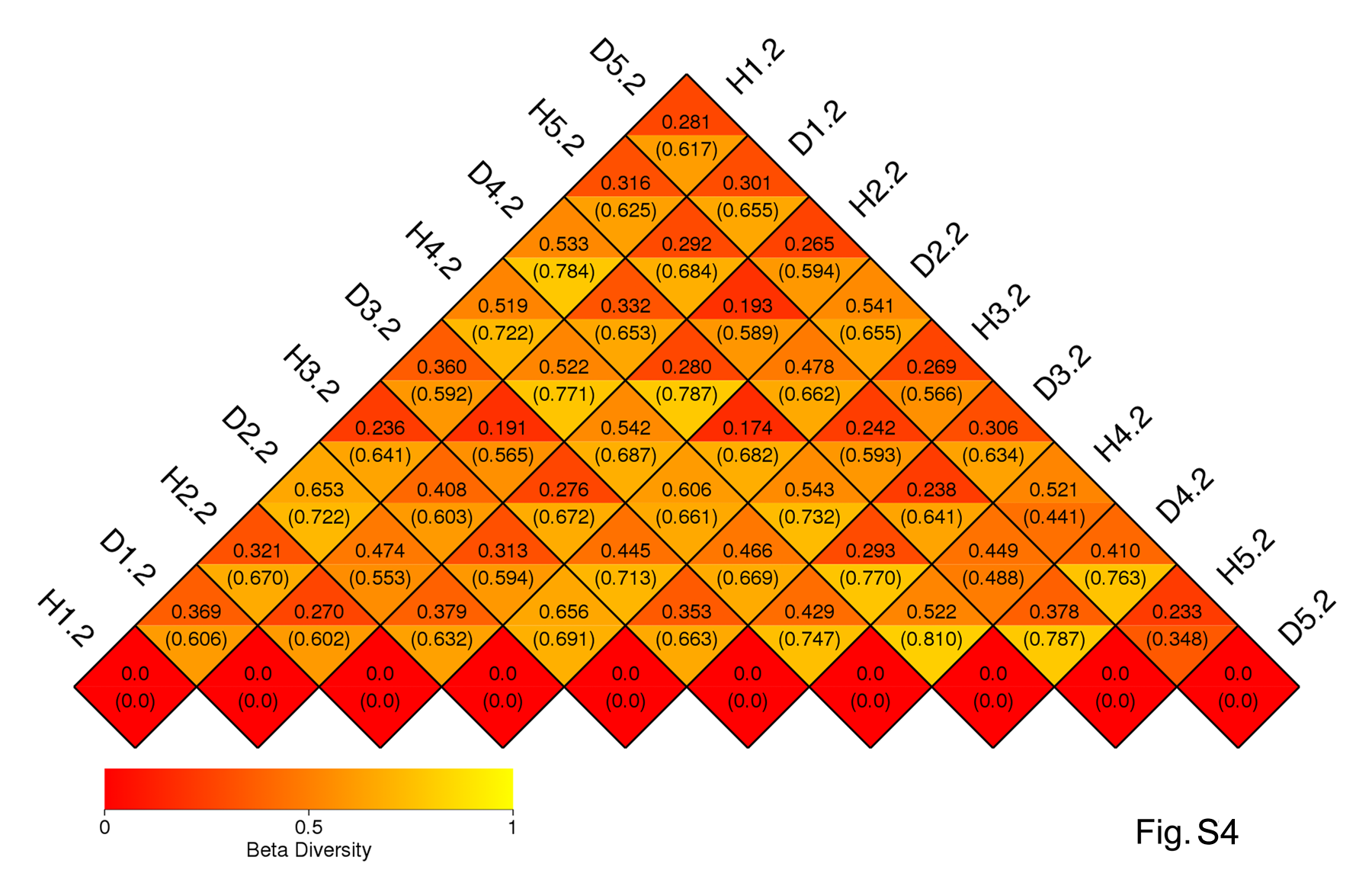

### Fig.S5

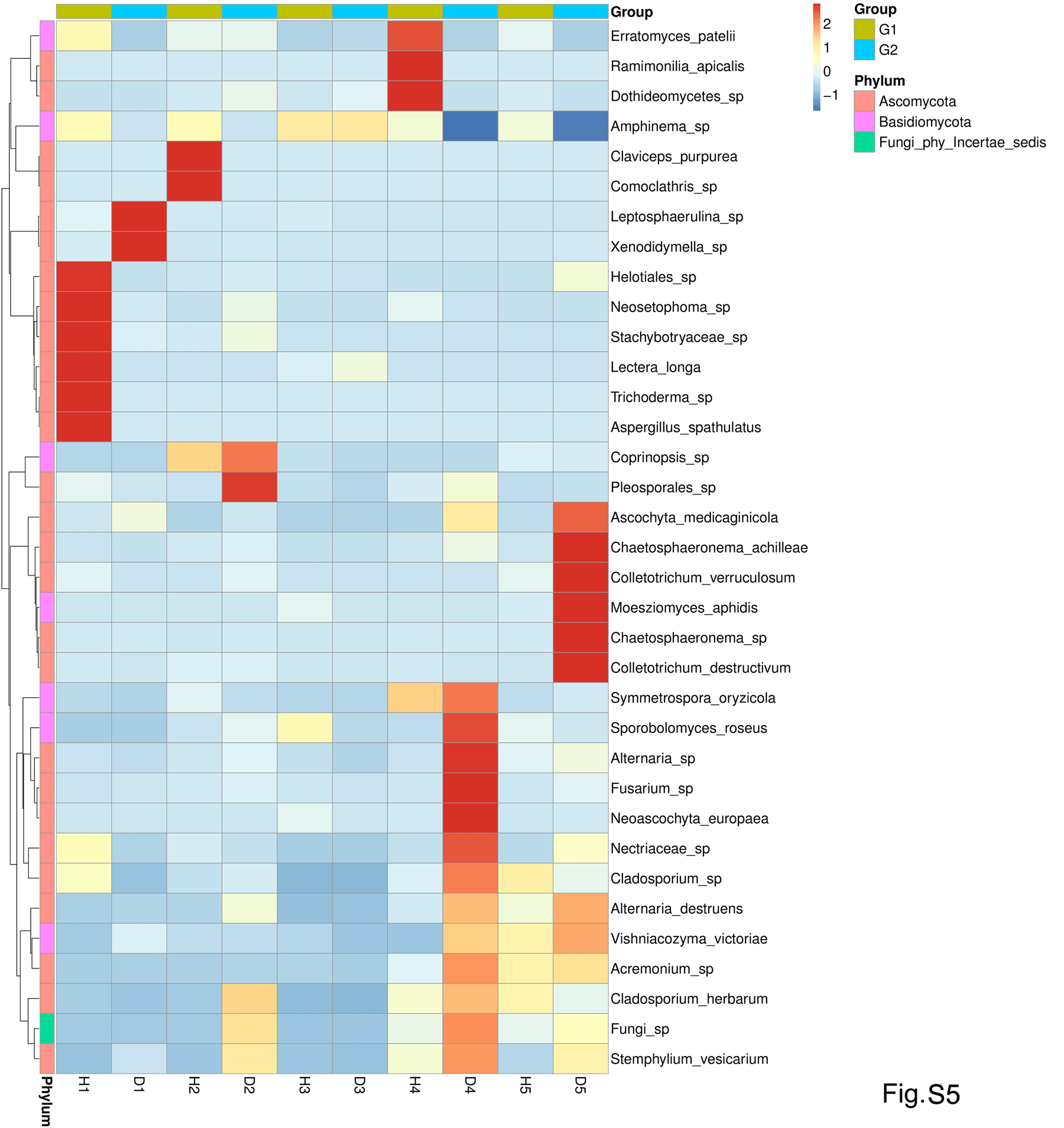

### Fig.S6

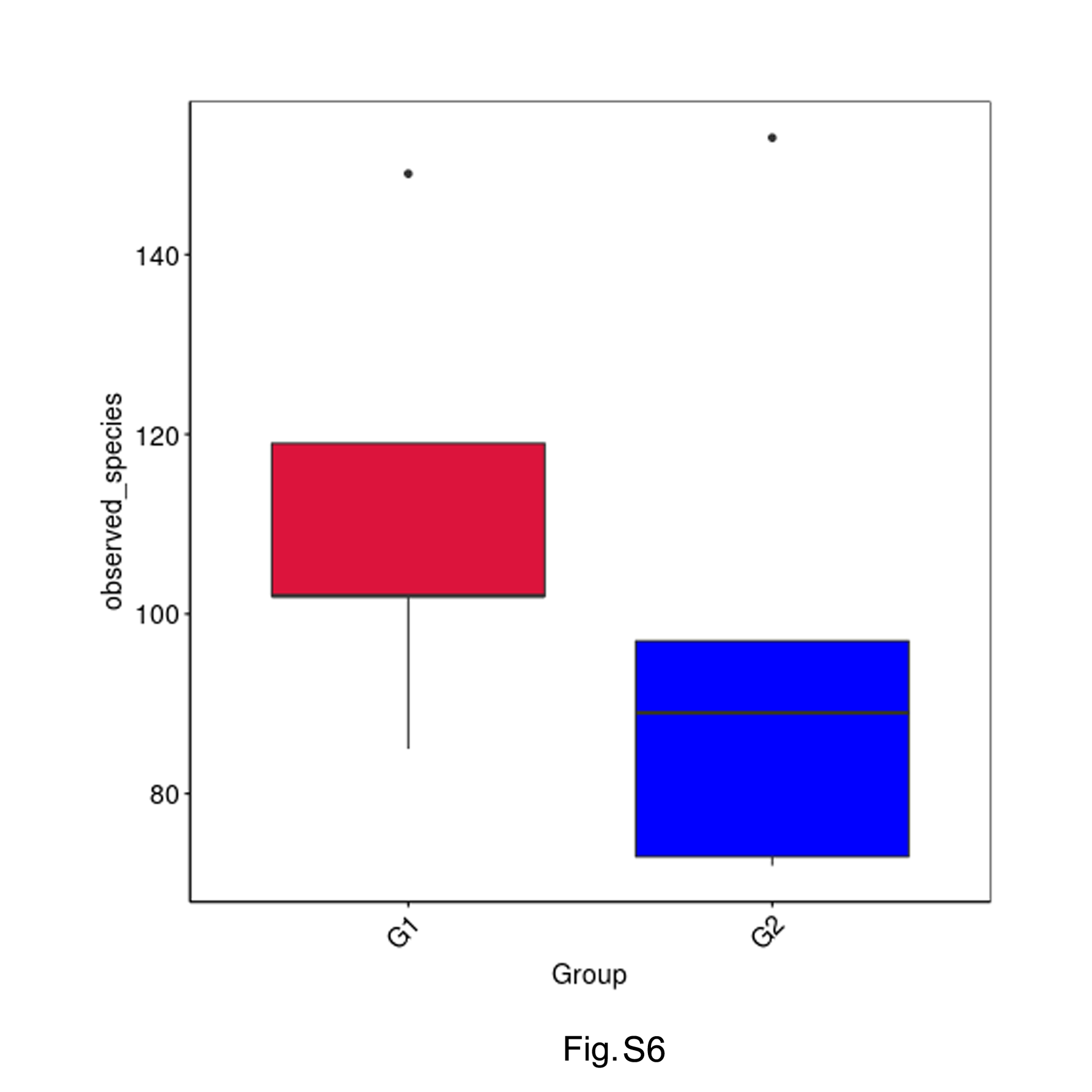

### Fig.S7

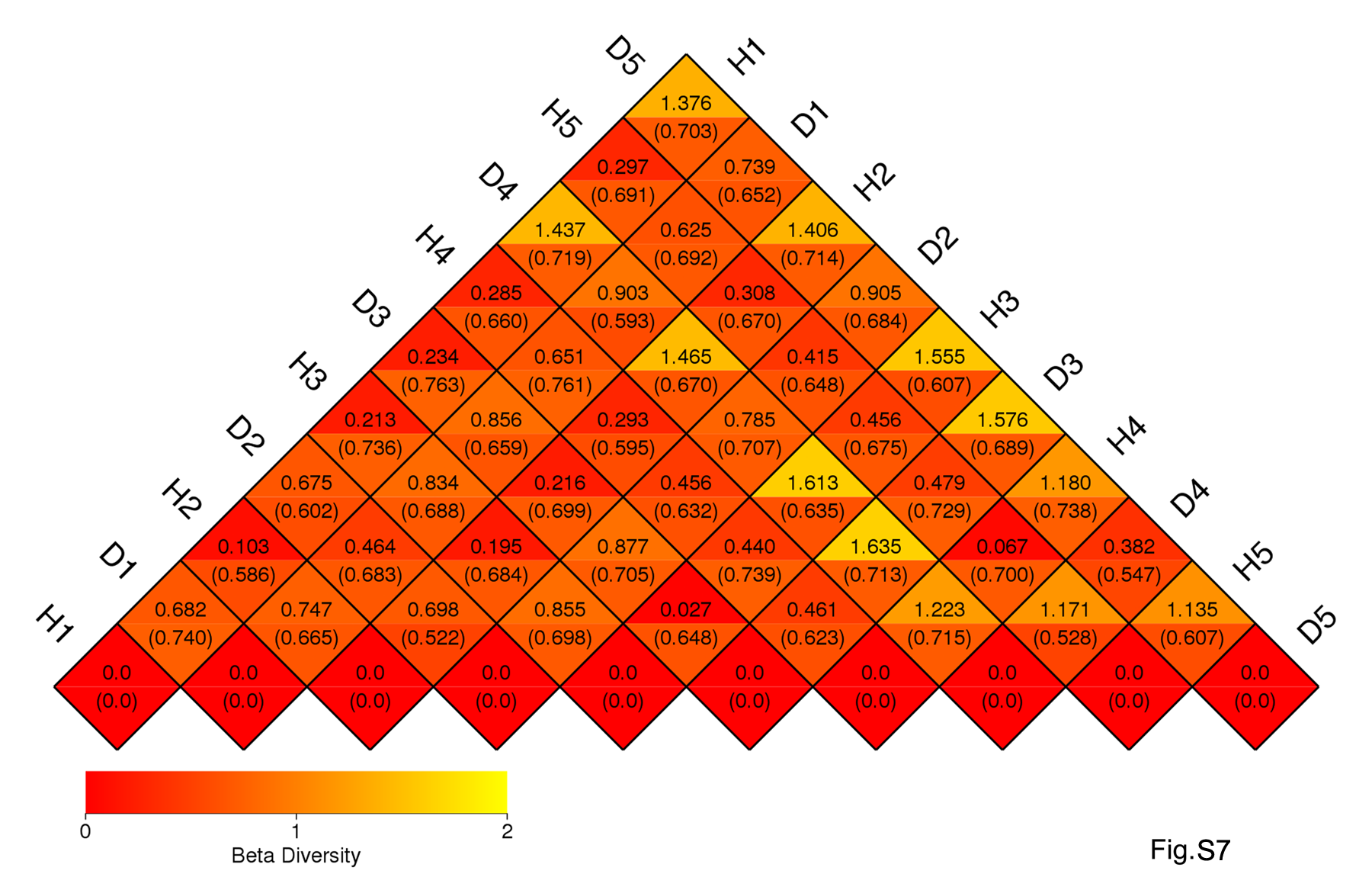

### Fig.S8

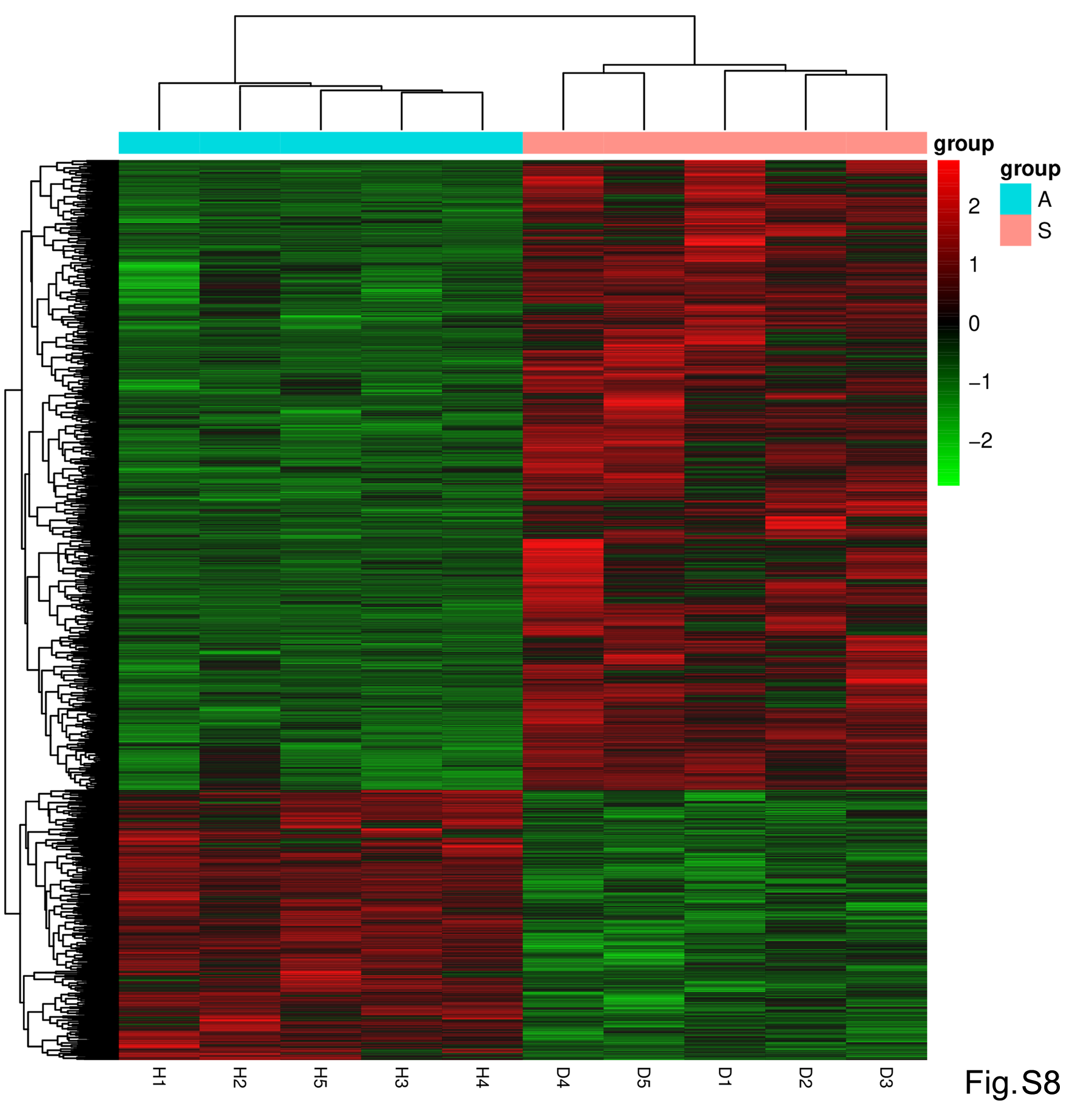

### Fig.S9

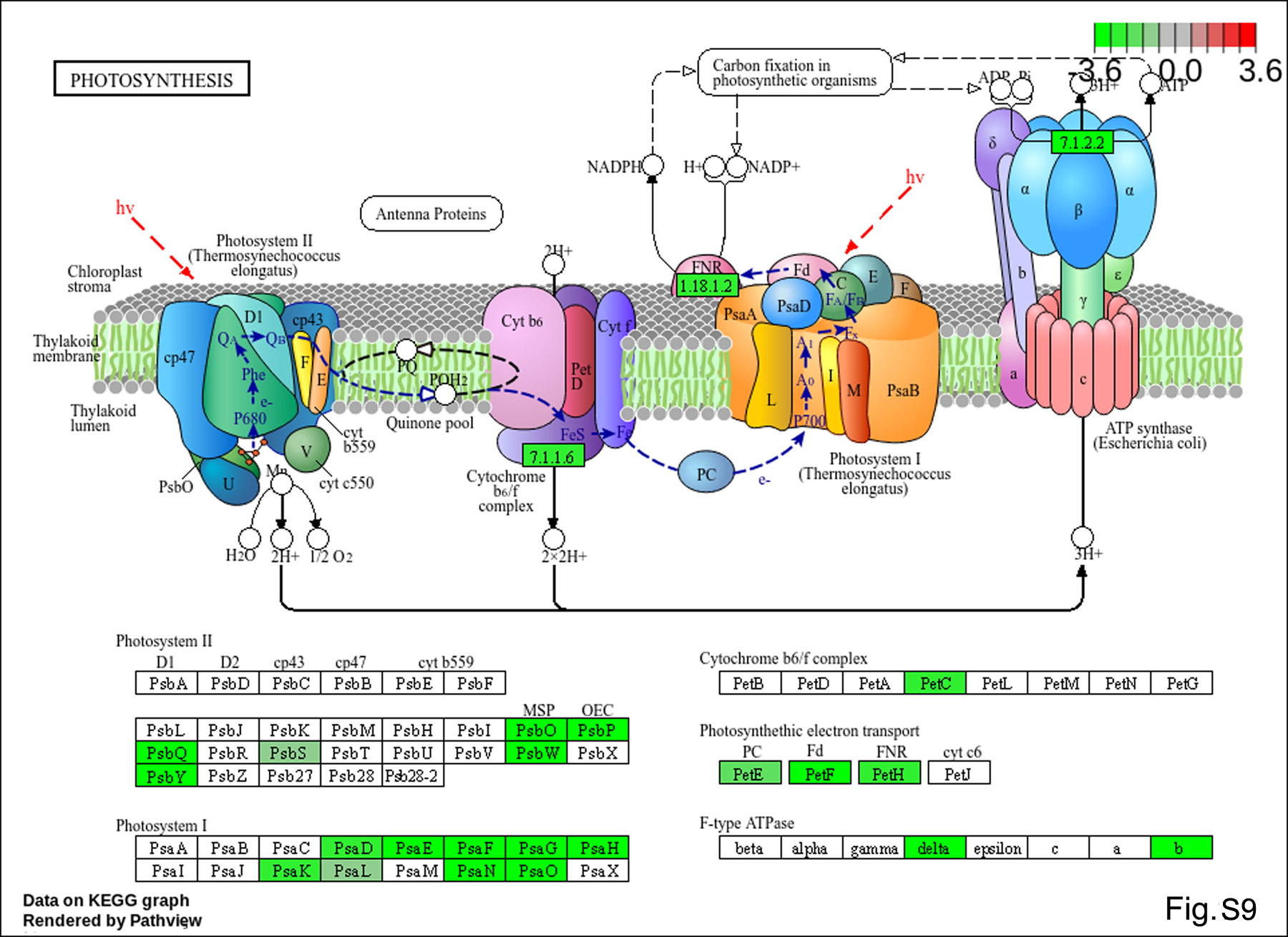

### Fig.S10

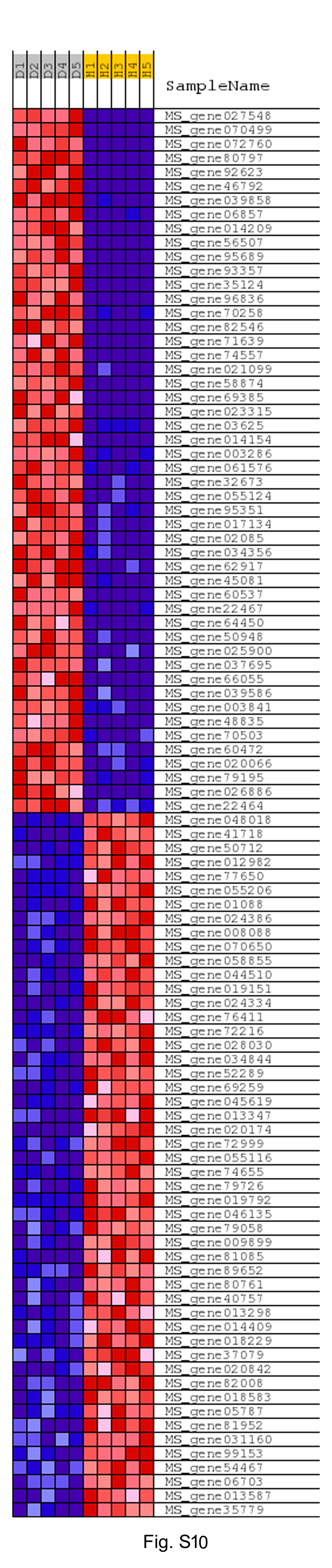

### Fig.S11

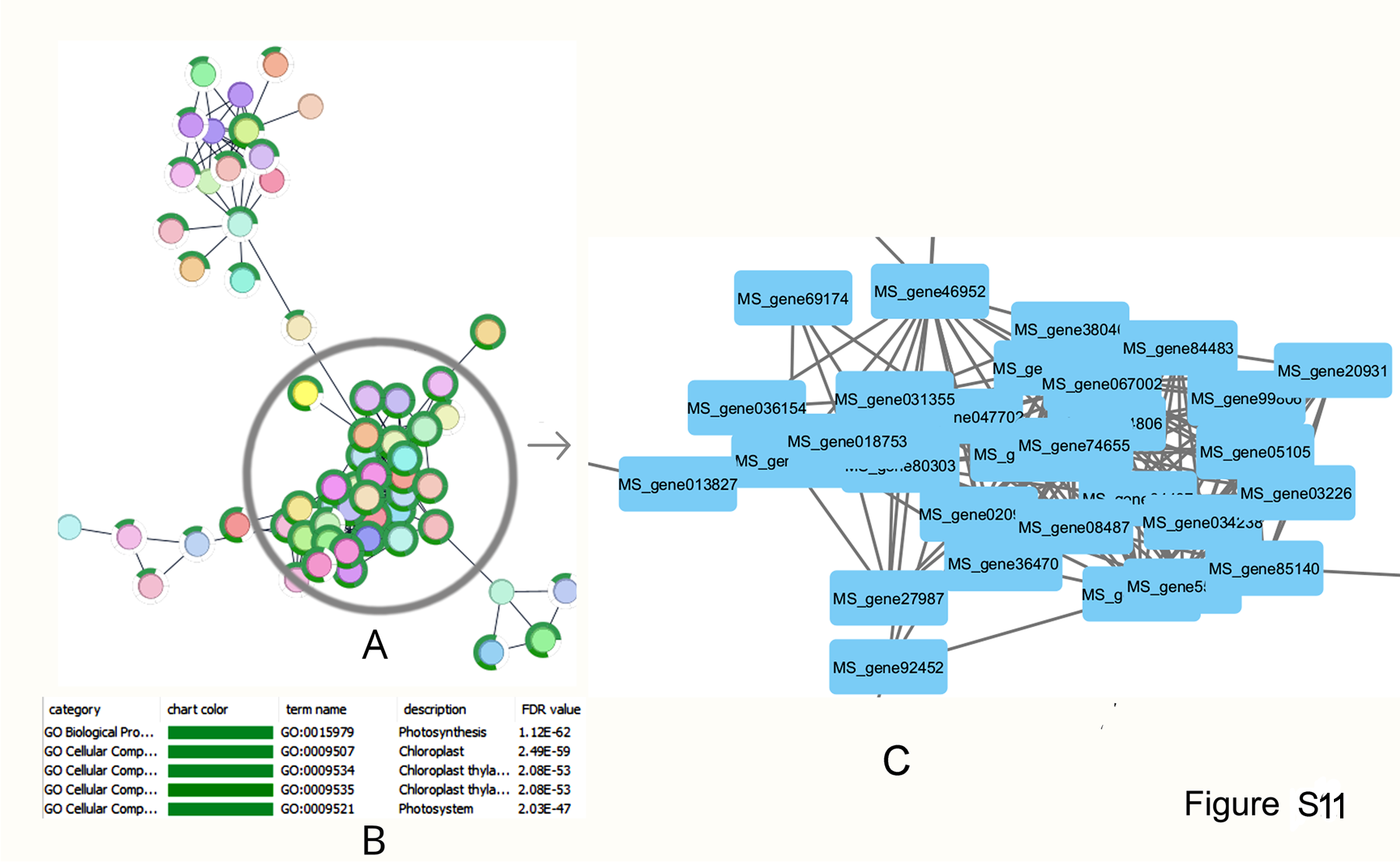
